# Profiling the microbial community structure and functional diversity of a dam-regulated river undergoing gravel bar restoration

**DOI:** 10.1101/2020.01.25.919381

**Authors:** Joeselle M. Serrana, Bin Li, Tetsuya Sumi, Yasuhiro Takemon, Kozo Watanabe

## Abstract

**Background:** River restoration efforts are expected to influence and change the diversity and functions of microbial communities following the recovery of habitat characteristics in the river ecosystem. The recreation or restoration of gravel bars in the Trinity River in California aims to rehabilitate the environmental heterogeneity downstream of the dam impounded channel. Here, we profiled the community composition, estimated diversity, and annotated putative metabolic functions of the sediment microbial communities to assess whether the construction and restoration of gravel bars in the Trinity River in California enhanced environmental heterogeneity, with the increase in the microbial beta diversity of these in-channel structures against the free-flowing reach of the main channel with comparison to its undisturbed tributaries.

**Results:** Microbial community composition of the free-flowing (i.e., no gravel bars) communities were relatively closer regardless of dam influence, whereas the Trinity River gravel bar and tributaries’ gravel bar communities were highly dissimilar. *Proteobacteria*, *Bacteroidetes*, and *Acidobacteria* were the highly abundant sediment microbial phyla on most sites, specifically in the Trinity River gravel bar communities. Putative functional annotation of microbial taxa revealed that chemoheterotrophy and aerobic chemoheterotrophy were the most prevalent microbial processes, with the Trinity River gravel bars having relatively higher representations. The considerably large abundance of heterotrophic taxa implies that gravel bars provide suitable areas for heterotrophic microorganisms with metabolic functions contributing to the net respiration in the river.

**Conclusions:** Our results provide supporting evidence on the positive impact of habitat restoration being conducted in the Trinity River with the non-dam influenced, undisturbed tributaries as the basis of comparison. Gravel bar recreation and restoration contributed to the increased microbial biodiversity through the restoration of environmental heterogeneity at the river scale. We provided valuable insights into the potential microbial processes in the sediment that might be contributing to the biogeochemical processes carried out by the microbial communities in the Trinity River. The significant positive correlation between the functional diversity of the identified microbial taxa and beta diversity suggests that differences in the detected metabolic functions were closely related to dissimilarities in community composition.

## Background

Dam construction and operation alters river hydrological regimes that directly influence various biogeochemical processes and ecosystem functions [1], which may induce community structure and functional homogenization in the dam-regulated river channel [2]. In particular, damming-induced changes are expected to influence beta diversity components of the downstream environment having huge implications for reservoir management and regional species diversity conservation [3]. Some restoration attempts on dam-impacted or sediment-limited lotic ecosystems aim to increase biogeochemical processing by integrating gravel augmentation to improve the hydrologic, geomorphic, and ecological processes in the river channel [4–5]. In-channel gravel features, e.g., ripples, bars, and meanders created by the addition of coarse sediments, provide environmental heterogeneity to rivers by diversifying the hydrodynamic exchange and residence time distribution [6] to promote the reestablishment of normative rates and magnitudes of physicochemical and biological processes [7].

In particular, gravel bars provide areas of increased biogeochemical activities due to the enforced hydrodynamic exchange [8–9] retaining organic matter filtered from surface waters in the hyporheic zone (i.e., the interface between surface and groundwater) for the utilization of river biota, and enhances nutrient cycling with consequent benefits to ecosystem metabolism [10]. The rise in environmental heterogeneity is expected to influence an increase in beta diversity, given that more heterogeneous conditions produce greater variation in species composition among localities within a regional unit [11–13]. Habitat heterogeneity in the form of spatial chemical and nutrient differences sufficiently drives beta diversity between microbial community composition in a floodplain [14]. Additionally, different taxonomic groups are expected to have different functional metabolic profiles [15]. Parallel taxonomic and functional diversity has been observed for microbial communities [16]. A study on stream microbial communities revealed consistent profile trends between taxonomic composition and functional profile in response to different land-use i.e., agriculture, wetland, and forested suggesting its influence on their contribution to various stream biogeochemical processes [17].

The taxonomic or functional diversity of microbial communities have been used as indicators of lotic ecosystem health due to their essential roles in organic carbon compounds and nutrient biogeochemical cycling [9, 18]. Changes in physical and biogeochemical factors directly affect microbial diversity and community composition, which inherently influence river ecosystem processes [18–19]. Many studies have identified various environmental factors such as particulate organic matter [20], water chemistry [21], and granulometric features [22] regulating microbial assemblage and activity in river ecosystems. Fine-scale sediment characteristics such as grain size, shape, composition, and heterogenic distribution derived from catchment-scale geological processes determine most of the physical and chemical processes in the hyporheic zone [21, 23]. For example, accumulation of organic matter and dissolved interstitial oxygen concentrations were reported to be influenced by sediment grain size [24], and an increase in surface area promoting higher microbial abundance due to fine sediment loading [25]. On the other hand, excessive sedimentation can have detrimental environmental effects on both the physical and biogeochemical processes in the hyporheic zone due to its effects on hydraulic conductivity and permeability between the surface channel and the nutrient supply. Chen et al. [26] observed reduced bacterial diversity and shifts in community composition in riparian sediments due to changes in the physicochemical properties of sediments as a result of damming. High loading of fine sediments clogs interstitial hyporheic spaces reducing porosity and water exchange between ground and surface water [27–29]. Alongside the translocation of fine particulate organic matter, sedimentation is expected to induce changes in microbial activity associated with shifts in biogeochemical processes (e.g., change from nitrification to denitrification) [21].

River restoration activities brought about by the restoration of habitat characteristics are expected to influence the diversity and composition of sediment microbial communities [26, 30]. Hence, microbial communities can be used as sensitive indicators of environmental changes to provide insights on the progression of aquatic habitat restoration [31]. Here, we profiled the community composition, estimated diversity, and annotated putative metabolic functions via next-generation sequencing of the 16S rRNA gene amplicon (V4 hypervariable region) of the sediment microbes of gravel bars along the main channel of a river undergoing restoration due to the impacts of dam impoundment. We also explored the association of microbial structure against different environmental variables. Our study represents the first attempt to explore the sediment microbiome of the Trinity River, California, and utilize microbial community data as a sensitive and integrative tool for the rapid assessment of the environmental impacts of restoration management in the channel. Comprehensive restoration programs for the Trinity River, which has been impounded by the Trinity and the Lewiston dams since 1964, employs ecological flow restoration and gravel augmentation for recreating instream gravel features. The Trinity River Restoration Program (TRRP) conducts coarse-sediment augmentation to recreate or rehabilitate gravel bars either via fluvial deposition of locally added sediments or by mechanical construction of gravel islands and bars [32–33]. Ock et al. [5] reported an increase in hyporheic exchange and suspended particulate organic matter retention from restored gravel bars resulting in thermal heterogeneity and food availability along the Trinity River channel.

We hypothesize that the construction and restoration of gravel bars in the Trinity River enhance environmental heterogeneity, in particular, the rehabilitation of fine-scale sediment characteristics, with the beneficial increase in microbial community composition and functional diversity in the gravel bars against the free-flowing reach of the main channel with comparison to its undisturbed tributaries. The objectives of the present study were therefore to (i) profile the microbial community structure of the sediments downstream of the dam-impounded Trinity River in comparison to non-dam impacted sites, (ii) determine how the gravel bar sediment communities differed from the non-in-channel microbial samples (sediments from free-flowing segments of the river) on both the Trinity River and its tributaries, (iii) identify indicator taxa and annotate the metabolic potential of these microbial communities, and (iv) classify the primary environmental factors influencing their composition, and determine its possible impact on the microbial metabolic potential of the restored in-channel gravel features.

## Results

### Variation in environmental variables

To assess the influence of dam-impoundment and presence of gravel bars on the sediment microbiomes of the Trinity River and its tributaries, we performed a paired sampling scheme with three spatial hierarchies, i.e., “River” [2 groups: dam-influenced (Trinity River, T) vs. non-dam influenced rivers (Reference tributaries, R)], (ii) “Reach” [2 groups: (gravel bar, B vs. free-flowing segment, F] and (iii) “Point” [4 groups: up- (UZ) and down-welling (DZ) points of the gravel bars, or up- (US) and downstream (DS) points of the free-flowing segments with comparable distances], and their combinations (iv) “River × Reach” (4 groups: T-B, T-F, R-B, and R-F), and (v) “River × Reach × Point” (8 groups: T-BDZ, T-BUZ, T-FUS, T-FDS, R-BDZ, R-BUZ, R-FUS, and R-FDS). Various environmental and water quality parameters were assessed for all twenty-four sampling points (Fig. S1, Table S1). Homogeneity of multivariate dispersion (PERMDISP) was used to test if differences in environmental conditions existed among the samples based on different categorical factors (See Table S2 for the PERMDISP *F*-statistics and *P*-values). pH and total suspended solids (TSS) showed significant differences in the environmental heterogeneity for “Reach” (2 levels - between the B and F samples) and “River × Reach” (4 levels-between the T-B, T-F, R-B, and R-F samples). Ammonium nitrogen (NH_4_-N), ash-free dry mass (AFDM), and nitrate nitrogen (N0_3_-N) were also significantly different among the “River × Reach × Point” (8 levels - between the T-BDZ, T-BUZ, T-FUS, T-FDS, R-BDZ, R-BUZ, R-FUS, and R-FDS samples). Electric conductivity (EC) did not only show a significant difference for both the “River × Reach” and “River × Reach × Point” categorical factors but also for “River” (between the T and R samples).

### Taxonomic identification of 16s rRNA reads

A total of 6,439,111 raw sequences were obtained from the twenty-four sediment samples. After quality filtering, denoising, and removal of chimeric sequences, clustering and chimera checking produced 2,855 amplicon sequence variants (ASVs) (Table S3). From this, 2,000 ASVs had taxonomic assignment (including unranked taxa). Two non-bacterial ASVs (seven reads) were excluded, and the remaining 1,998 ASVs were assigned to 152 ranked and 173 unranked (e.g., “unclassified *Burkholderiaceae*”) bacterial genera belonging to 231 families, 151 orders, and 61 classes, under 27 phyla (including “unclassified bacteria” assignment). An overview of the microbial taxonomic assignment is provided as a browsable Krona chart (Additional file 1).

At the genus-level assignment, “unclassified *Burkholderiaceae*” and “unclassified *Rhodobacteraceae*” under *Proteobacteria* were the most dominant, with 11.4% and 6.65% read abundances, followed by *Methylotenera*, *Sphingorhabdus*, *Flavobacterium*, *Nitrospira*, and *Fluviicola* with more than 3% of the total read abundance. A heatmap representation of the genus-level assignment for each of the twenty-four communities is presented in Figure S2. At the class level, *Alphaproteobacteria*, *Gammaproteobacteria*, *Bacteroidia*, *Blastocatellia* (Subgroup 4), *Deltaproteobacteria*, *Nitrospira, Verrucomicrobiae*, *Acidobacteria* (Subgroup 6), *Planctomycetes* OM190, and *Planctomycetacia* were dominant with at least 1% from the total reads. At the phylum level, *Proteobacteria* was the most dominant (59.3%), followed by *Bacteroidetes* (18.1%), *Acidobacteria* (7.2%), *Nitrospirae* (4.2%), *Verrucomicrobia* (3.8%), and *Planctomycetes* (3.6%) which accounted for 96.2% of the total bacterial sequences, and appears on most of the sampling points. Also, *Gemmatimonadetes*, *Cyanobacteria*, *Elusimicrobia*, and *Chloroflexi* were relatively abundant (with more than 0.4% of the sequences) with a large sample to sample variation. A heatmap representation of the phylum-level assignment for each of the twenty-four communities is presented in Figure S3.

### Microbial richness and diversity measures

Alpha diversity indicators were calculated (Table S4). The richness indices of the 24 sampling points for Chao1 and Fisher’s alpha metrics approximately ranged from 3 to 509 and 0.37 to 129, with gravel bar points B6UZ, B6DZ, B8UZ and B8DZ, and free-flowing points F1DS and F1US represented with the lowest values. Similarly, the microbial diversity indices, i.e., Simpson and Shannon, were relatively low for the down-welling points of B6DZ and B8DZ. The mean alpha diversity values for the groups on each of the categorical factors are presented in Figure S4. Mean values of all the alpha diversity indices, including the observed ASVs (richness estimate), were relatively higher for the Trinity River in comparison to the reference creeks, and the gravel bar reaches in comparison to free-flowing reaches. In terms of the grouping based on “River × Reach × Point”, both the down- (T-BDZ) and up-welling (T-BUZ) points of the dam-impacted bar communities have relatively higher Chao1 values 73.76 and 146.19 and Fisher’s alpha values 17.10 and 36.23, respectively. T-BDZ and T-BUZ also have relatively high values, with the addition of the up-welling points of the non-dam influenced gravel bar reaches (R-BUZ). However, only the Simpson (*t-value* = 1.75, *P* = 0.093) and Shannon (*t-value* = 1.95, *P* = 0.064) values of the categorical factor “River” showed statistical significance at *P* = 0.10 (Table 1).

**Table 1.** Significance test of alpha diversity estimates at different categorical factors.

| Categorical Factor | Observed ASVs |  | Chao1 |  | Fisher alpha |  | Simpson |  | Shannon |  |
| --- | --- | --- | --- | --- | --- | --- | --- | --- | --- | --- |
|  | *t-value* | *P* | *t-value* | *P* | *t-value* | *P* | *t-value* | *P* | *t-value* | *P* |
| (Student's t-test) |  |  |  |  |  |  |  |  |  |  |
| River | 1.610 | 0.120 | 1.420 | 0.169 | 1.430 | 0.170 | 1.750 | **0.093** | 1.950 | **0.064** |
| Reach | 1.449 | 0.161 | 1.308 | 0.204 | 1.347 | 1.192 | 1.510 | 0.145 | 1.833 | 0.803 |
| (1-way ANOVA) | *F* | *P* | *F* | *P* | *F* | *P* | *F* | *P* | *F* | *P* |
| Point | 0.900 | 0.459 | 0.850 | 0.483 | 0.900 | 0.459 | 1.480 | 0.250 | 1.330 | 0.293 |
| River × Reach | 0.730 | 0.064 | 0.639 | 0.718 | 0.658 | 0.704 | 0.918 | 0.518 | 0.906 | 0.526 |
| River × Reach × Point | 1.090 | 0.370 | 1.270 | 0.312 | 1.300 | 0.302 | 1.410 | 0.269 | 2.140 | 0.127 |

Student's t-test for factors with 2-levels, and 1-way ANOVA for independent samples for factors with more than 2: "River" (2 groups: Trinity River – dam- influenced, Reference River – non dam-influenced); "Reach" (2 groups: gravel bars, free-flowing segment); "Point" (4 groups: up- and down-welling, and up- and down-stream); "River × Reach" and "River × Reach × Point" with 4 and 8 groups, respectively. *F*: F-statistic; *P*: p-value. Significant effects and interactions are highlighted in bold (significance value < 0.1).

### Differences in microbial community structure between categorical factors

Plots from principal coordinates analysis (PCoA) of each categorical factor were presented in Figure 1 (A-E). PCoA revealed that the microbial communities were relatively distinct between the Trinity River and its tributaries (by “River”; Fig.1a), and between the gravel bars and the free-flowing reach (by “Reach”; Fig.1b), with the former having a small overlap. In accordance, the samples categorized by “Point” (Fig.1c) revealed UZ and DZ of the gravel bars closely clustered together, and the DS and US of the free-flowing reach separated from the latter, with a relatively small overlap. PCoA on “River × Reach” (Fig.1d) showed that the free-flowing reaches were clustered relatively close regardless of the river, indicating closely similar community composition in the samples from the non-gravel bar communities uninfluenced by the presence or absence of dam-impact. In contrast, both gravel bar communities from the Trinity River and the reference creeks were separately clustered relatively far against the free-flowing reaches and were clustered on separate corners of the ordination plot, suggesting a high dissimilarity in the community composition between the gravel bar reaches of the two groups. Fig. 1e also presents the PCoA plot of the “River × Reach × Point” category. The non-metric multidimensional scaling (NMDS) ordination plots based on Bray-Curtis distance reflects similar ordination as their PCoA counterparts (Fig. S5).

**Fig. 1.**
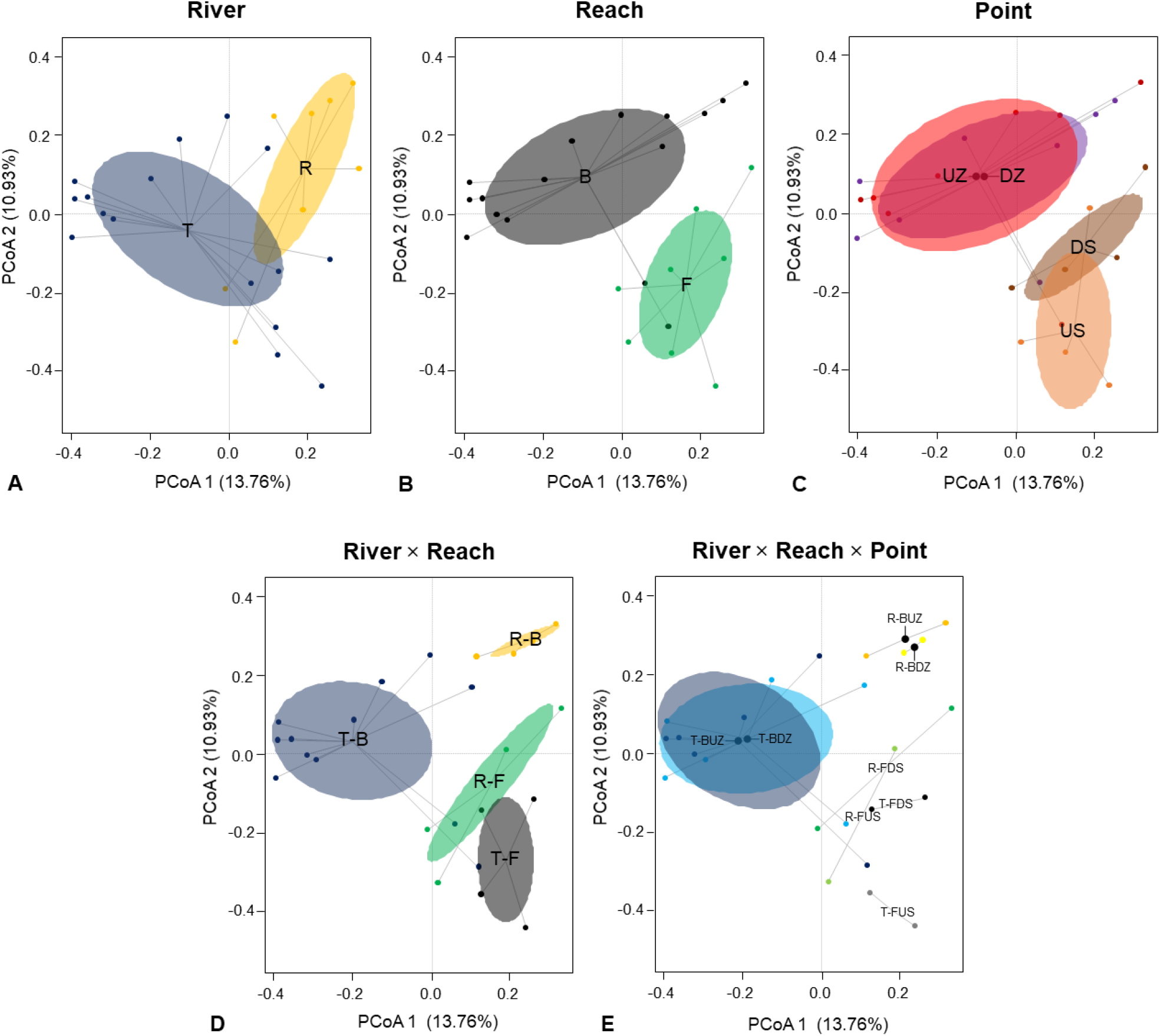
Effect of categorical factors on the beta diversity of microbial communities at the genus-level represented via Principal coordinates analysis (PCoA) based on Bray-Curtis distance from the PERMDISP2 analysis (*betadisper* function). Groupings based on the categorical factors are presented with ellipses (95% confidence) representing the standard error around the centroid. **a** River: “T” stands for Trinity River (dam-influenced); “R” for reference creek (no dam-influence). **b** Reach: “B” for gravel bar; “F” for the free-flowing segment of the river. **c** Point: “UZ” for up- and “DZ” for down-welling zones of the gravel bar sites; “US” for up- and “DS” for downstream sampling points of the free-flowing river segments, and their combinations: **d** “River × Reach”, and **e** “River × Reach × Point”.

The analysis of similarity (ANOSIM) test revealed that microbial communities were significantly differentiated across all the categorical groups and combinations, with “River” having the lowest *R*-value (*R*_ANOSIM_ = 0.18, *P*_ANOSIM_ = 0.017). A value of *R* = 0 indicates equal similarity between and within samples, while a value of *R* = 1 indicates higher similarities within than between categorical groups. On the other hand, the permutational multivariate analysis of variance (PERMANOVA) revealed that “River” (*R^2^*_PERMANOVA_ = 0.08, *P*_PERMANOVA_ = 0.0015) and “Reach” (*R^2^*_PERMANOVA_ = 0.09, *P*_PERMANOVA_ < 0.001) significantly influenced microbial community dissimilarity across all samples at *P* = 0.05, while “River × Reach” (*R^2^*_PERMANOVA_ = 0.05, *P*_PERMANOVA_ = 0.0806)was not significant at *P* = 0.05 (Fig. S6; Table 2). A separate test of homogeneity of dispersion via Permutational Analyses of Multivariate Dispersions (PERMDISP) followed by an analysis of variance (ANOVA) test showed significant differences in the microbial community dispersion (within-group variation in beta-diversity) among all of the categorical groups and combinations (i.e., “River × Reach” and “River × Reach × Point”) except for “River” (*F*_PERMDISP_ = 1.86, *P*_PERMDISP_ = 0.193) (Fig. S6; Table 2).

**Table 2.** Statistical difference in beta diversity.

| Categorical Factor | ANOSIM |  | PERMANOVA |  |  |  |  | PERMDISP2 |  |  |  |
| --- | --- | --- | --- | --- | --- | --- | --- | --- | --- | --- | --- |
|  | *R* | *P* | Df | SS | *R*<sup>2</sup> | *F* | *P* | Df | SS | *F* | *P* |
| River | 0.18 | **0.0170** | 1 | 0.78 | 0.08 | 2.00 | **0.0015** | 1 | 0.01 | 1.86 | 0.1865 |
| Reach | 0.21 | **0.0030** | 1 | 0.92 | 0.09 | 2.35 | **0.0001** | 1 | 0.01 | 4.71 | **0.0411** |
| Point | 0.21 | **0.0100** | 2 | 0.90 | 0.09 | 1.15 | 0.1820 | 3 | 0.04 | 2.85 | 0.0632 |
| River × Reach | 0.35 | **0.0010** | 1 | 0.53 | 0.05 | 1.36 | 0.0806 | 3 | 0.05 | 2.69 | 0.0738 |
| River × Reach × Point | 0.32 | **0.0030** | 2 | 0.62 | 0.06 | 0.80 | 0.9135 | 7 | 0.15 | 5.65 | **0.0020** |

ANOSIM, PERMANOVA and PERMDISP2 analysis to test the effect of "River", "Reach", and "Point" on the microbial communities at the genus level. "River" (2 groups: Trinity River – dam-influenced, Reference River – non-dam-influenced); "Reach" (2 groups: gravel bars, free-flowing segment); "Point" (4 groups: up- and down-welling, and up- and down-stream). Df: degrees of freedom; SS: sum of squares; *Pseudo-F*: F value by permutation; *P*: p-value. Significant effects and interactions are highlighted in bold (significance value < 0.05).

### Indicator taxa associated with each categorical factor

Pairwise taxonomic comparison heat trees based on ASV abundance (≥0.10 total read abundance) for the “River × Reach” categories were generated via *metacoder* (Fig. 2) to identify the taxa influencing the similarity and differences between the groups. T-B was relatively more represented by *Alphaproteobacteria* and *Gammaproteobacteria* of the phylum *Proteobacteria*, and the genus *Flavobacterium* (*Bacteroidetes*) in comparison to the tributary sites i.e., R-B and R-F. Similarly, T-B was relatively different from T-F on the taxa as mentioned above except for *Gammaproteobacteria* which was relatively more abundant in the latter, specifically representatives from the family *Burkholderiaceae*. Relative to the gravel bar reaches, both T-F and RF have abundant representatives of the family *Blastocatellacea*. A figure including by “Reach” taxonomic comparison for a clearer view of the taxonomic tree is provided in the supplementary information (Fig. S7).

**Fig. 2.**
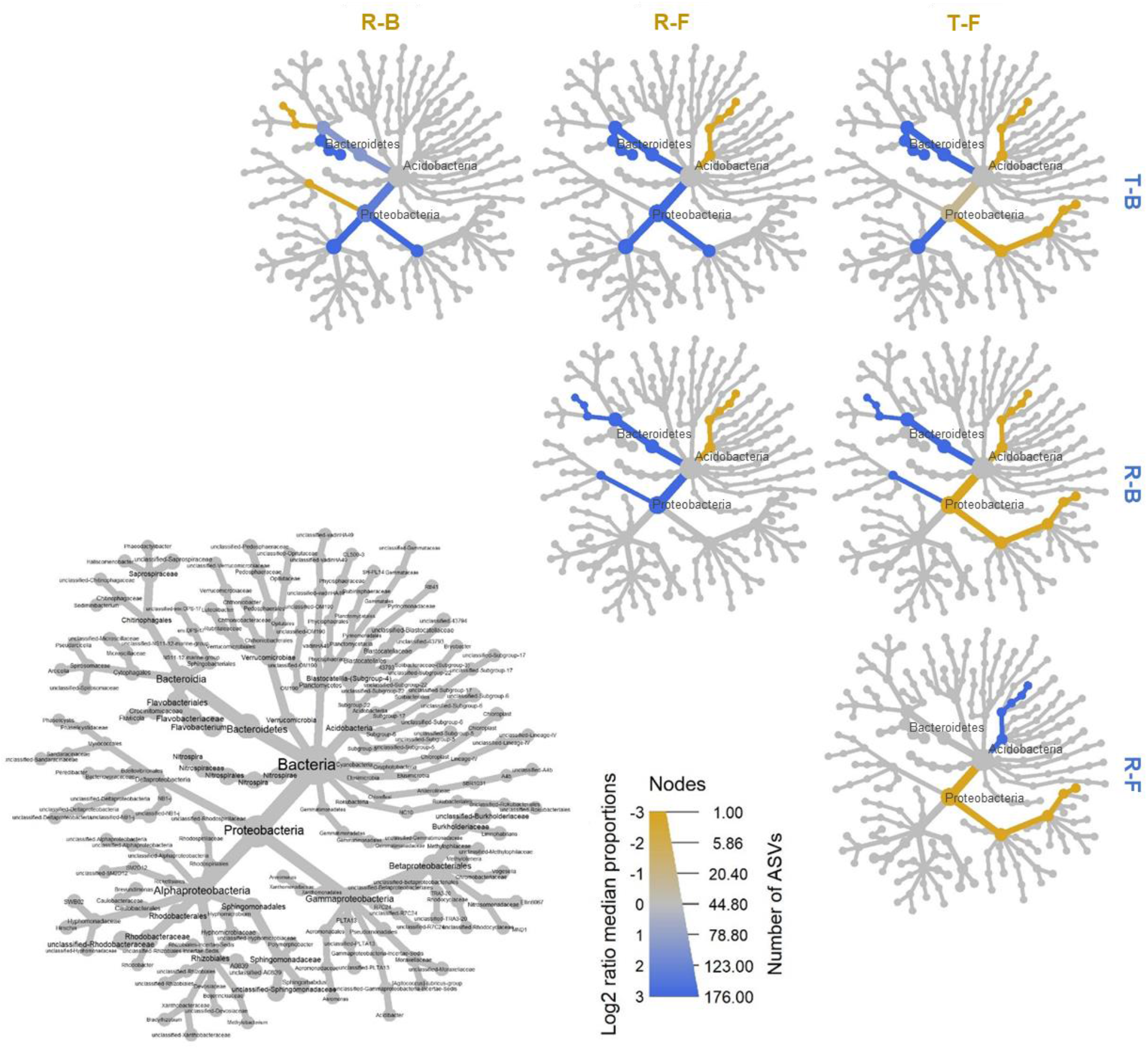
Pairwise taxonomic comparisons (ASVs with relative read abundance >0.10) of the “River × Reach” categories. The grey tree on the lower left functions as the key for the unlabeled trees. Each of the smaller trees represent a comparison between “River × Reach” categories in the column and rows. Taxon colored in yellow are relatively more abundant (in numbers of ASVs) in the category in the column and a taxon colored blue is more abundant in the category of the row. Legend: width indicates number of ASVs at a given taxonomic rank, and color indicates relative differences in log2 (number of ASVs). A figure including “Reach” taxonomic comparison for a clearer view of the taxonomic tree is provided in the supplementary information (Additional file 2: Fig. S7).

Indicator taxa analysis revealed distinct taxa associated with each group of samples categorized based on dam-influence, or reach type. Indicator taxa represent taxa that are abundant in a group but rare or absent in other groups. Sixteen genera (including unranked taxa) out of 325 were identified as significant indicator taxa (*P* = 0.10) based on the “River × Reach” category whose abundances may be reflective of particular environmental parameters (Table S5). *Flavobacterium*, *Sediminibacterium*, and unclassified *Rhodobacteraceae* were identified indicators of the T-B category. *Pirellula*, *Deefgea*, and three unclassified families, i.e., *Gemmataceae*, *Microscillaceae*, and *Sporichthyaceae*, were indicators of the R-B category. The T-F category has indicator taxa from *Haemophilus*, SM1A02 (*Phycisphaeraceae*), two unclassified families, i.e., *Burkholderiaceae*, and *Rhodocyclaceae*, and a group of unclassified *Gammaproteobacteria*. Only the genus *Acinetobacter* was detected as indicator taxa for the R-F category. Additionally, unclassified *Betaproteobacteriales* were indicator taxa for the free-flowing reaches (i.e., T-F and R-F) and a group of unclassified *Blastocatellaceae* for the R-B, T-F and R-F categories.

### Correlation of microbial community distribution and environmental variables

Bray-Curtis distance-based redundancy analysis (dbRDA) was performed to evaluate the relationship between the taxonomic compositions of microbial communities and the environmental variables (Fig. 3). All seven environmental parameters assessed in this study have less than or equal to a variance inflation factor (VIF) score of 3. The global dbRDA model was statistically significant following a permutational ANOVA test (*F* = 1.33, *P* = 0.0024). The first and second axis of the dbRDA plot explained 10.5% and 17.7% of the variation in the data, respectively. The first axis was statistically significant (*F* = 2.66, *P* = 0.005), and was negatively correlated with NO_3_-N and TSS, but positively correlated with DO, EC, NH_4_-N, pH, and AFDM. However, only the last two environmental variables were significantly explained following a permutational ANOVA test (pH: *F* = 1.43, *P* = 0.045; AFDM: *F* = 1.33, *P* = 0.092). Free-flowing reach (i.e., T-F and R-F) samples were associated with high DO, and the non-dam influenced (i.e., R-F and R-B) samples were associated with high pH and AFDM, whereas T-B was mostly associated with TSS and NO_3_-N concentrations.

**Fig. 3.**
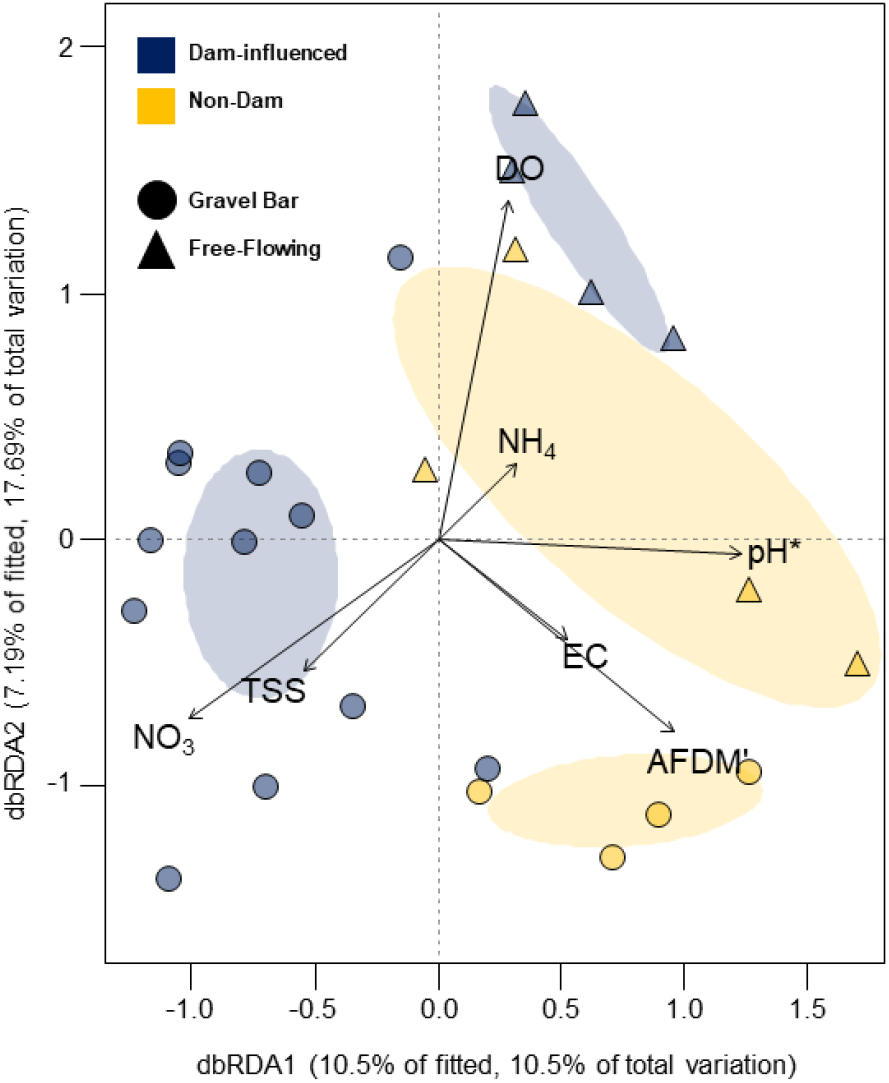
Distance-based redundancy analysis (dbRDA) showing the relationship between the microbial community structure (at the genus-level) and the environmental parameters. Grouping based on the categorical factor “River × Reach” is presented with ellipses (95% confidence) representing the standard error around the centroid. Significance code: ^‘**’^ associated with a variable at *P* < 0.01, ^‘*’^ at *P* < 0.05 and ^‘′’^ at *P* < 0.1.

### Estimation of microbial metabolism function

A total of 458 out of 1,998 ASVs (22.9%) were assigned to at least one functional group, with 38 from 90 functional groups represented (i.e., associated with at least one record). ASVs without any functional annotation (or leftovers; 77.1% of the ASVs) were excluded from the following analysis. Five sampling points, i.e., B6DZ, B8DZ, B8UZ, F1US, and F1DS, did not have functional annotations. The cumulative abundance of the ASVs assigned to each functional group was used to calculate the relative abundance in each sample after normalizing the cumulative abundances of the ASVs associated with at least one function (Fig. S8). Chemoheterotrophy was the most commonly annotated function for most microbial communities except for the samples without any annotated functions and B5DZ with only a predatory or exoparasitic group match. Aerobic chemoheterotrophy was also prominent for most of the gravel bar samples. Grouping the functional groups based on “River × Reach” revealed T-B has 34 represented functional groups, followed by T-F and R-F with fourteen and thirteen respectively (Fig. 4a). In addition to chemoheterotrophy, aerobic chemoheterotrophy and nitrate reduction were the most prominent metabolic functions in the T-F samples. Both T-F and T-B have methanotrophy, hydrocarbon degradation, and methylotrophy functions. The R-F samples were the only reaches with dark sulfide oxidation, and dark oxidation of sulfur compounds metabolic functions. Whereas R-B only matched with ten functional groups, with chemoheterotrophy, aerobic chemoheterotrophy, predatory, and cellulolysis being highly represented.

**Fig. 4.**
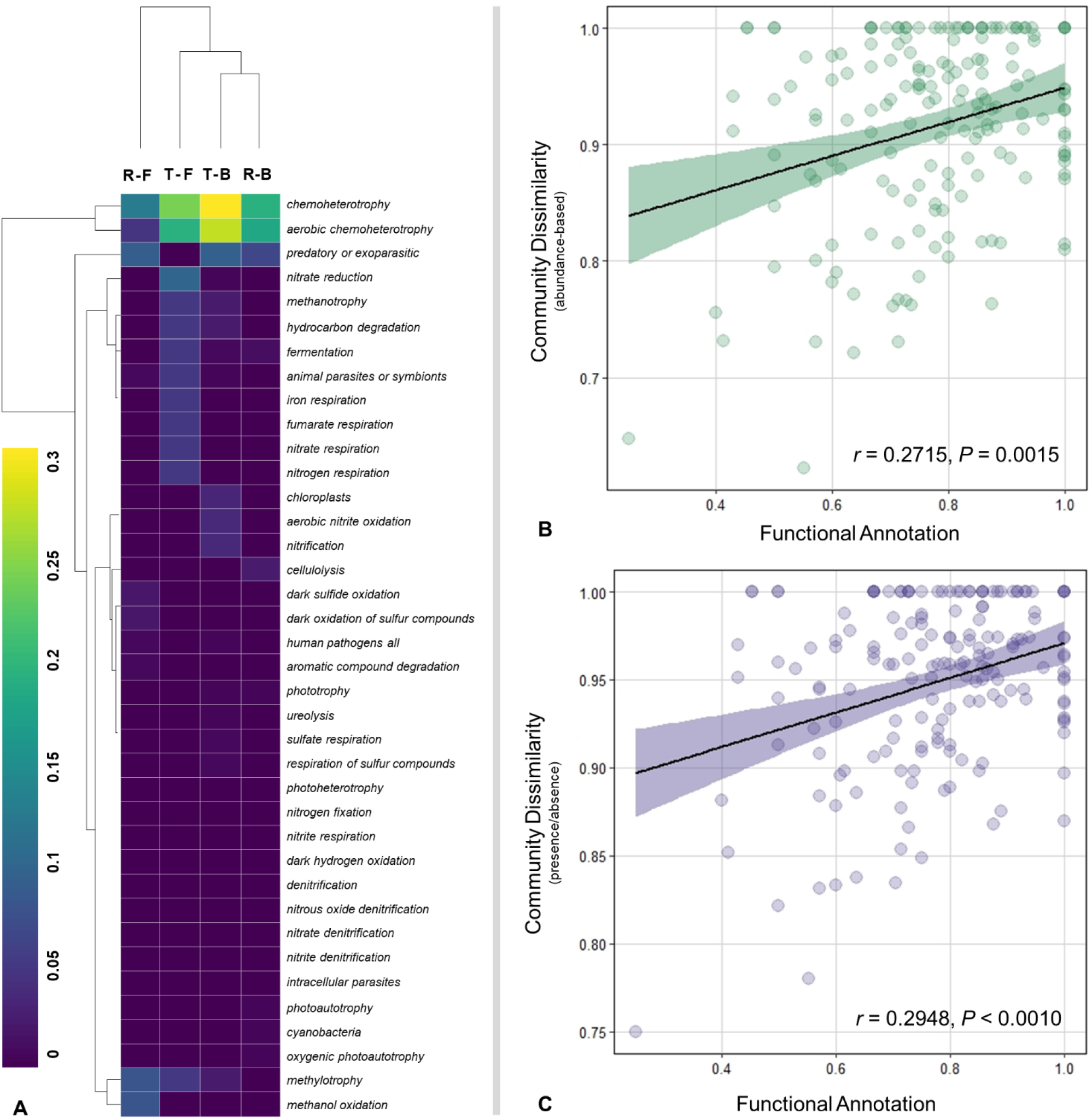
Functional annotation of the microbial communities. **a** The relative abundance of the predicted functional groups based on the categorical factor “River × Reach”. Double hierarchical cluster-based on Euclidean distance. Mantel test showing the relationship between community dissimilarity and functional annotation. **b** Beta diversity based on Bray-Curtis distance (read abundance) at the genus-level and functional annotation (Jaccard distance for presence/absence data). **c** Beta diversity based on Jaccard distance (incidence, presence/absence) at the genus-level and functional annotation (Jaccard distance). The figure denotes 95% confidence interval.

The functional dissimilarity calculated based on Jaccard distance was significantly correlated with the community dissimilarity estimated at the genus-level (Bray-Curtis distance) for both the Mantel test (*r* = 0.2715, *P* = 0.0015) (Fig. 4b) and the Procrustes test (*m^2^* = 0.2801, correlation in a symmetric Procrustes rotation = 0.8485, *P* < 0.001). Similarly, functional dissimilarity showed significant correlation with the incidence-based community dissimilarity estimated at the genus-level (Jaccard distance) for both the Mantel test (*r* = 0.2948, *P* = 0.0008) (Fig. 4c) and the Procrustes test (*m^2^* = 0.2734, correlation in a symmetric Procrustes rotation = 0.8524, *P* < 0.001).

## Discussion

The comprehensive restoration programs for the Trinity River implementing gravel augmentation and ecological flow restoration for recreating instream gravel features [4–5] are expected to influence the diversity and composition of sediment microbial communities in accordance to changes attributed to the restoration of habitat characteristics [26, 30] in the dam-impacted channel. Our results revealed significant compositional differences between the gravel bars and the free-flowing communities (by “Reach”; Fig. 1B) for all the non-parametric analyses of beta diversity. Gravel bars and free-flowing reaches are expected to show the difference in community composition, given that the former supports higher microbial biomass and anaerobic conditions due to its partially submerged geomorphic structures in comparison to their fully submerged counterparts [34]. The clustering analyses (PCoA or NMDS) based on Bray-Curtis distance supports the clear separation between the T-B and the R-B communities clustered on separate corners of the ordination space, whereas the free-flowing reaches, i.e., T-F and R-F clustered relatively closer regardless of dam influence.

The observed difference between the gravel bar microbial communities can be attributed to their differences in bar geomorphology and sediment characteristics. Most of the gravel bars in the Trinity River were constructed via fluvial deposition of locally added sediments or by mechanical construction of gravel islands and bars [32], whereas the tributaries assessed were naturally formed gravel bars. Claret et al. [35] reported that gravel bars located in a stable area of the river have low transversal heterogeneity, enabling aerobic processes such as mineralization of organic matter and nitrification due to low fine sediment content and high oxygenation of interstitial water. Whereas, gravel bars in the degraded zones had spatially variable structures acting as a dissolved organic carbon sink and nitrate source due to the localized accumulation of fine sediment and hypoxic conditions promoting anaerobic processes such as denitrification. The restored gravel bars in the Trinity River have been reported to produce hydraulic gradients by creating bed topography, introduced highly permeable freshly-deposited coarse sediments, which increased channel complexity [5].

River restoration projects are conducted under the assumption that physical habitat restoration will increase biodiversity [36]. In particular, channel re-configuration by adding meanders and physical structures, e.g. artificial rifles and boulders, restore biodiversity by enhancing stream structural heterogeneity of stream channels [37]. Singer et al. [38] highlighted the importance of fine-scale streambed heterogeneity for microbial biodiversity and ecosystem functioning, with environmental homogenization and loss of habitats increasingly reduce biodiversity. Additionally, stream mesocosm experiments revealed that microbial beta diversity increased with habitat heterogeneity at the landscape scale [39]. Our results provide supporting evidence on the positive impact of habitat restoration being conducted in the Trinity River with the non-dam influenced, undisturbed tributaries as the basis of comparison. Construction or restoration of gravel bars contributed to the increased microbial community dissimilarity through the restoration of environmental heterogeneity at the river scale. This study is the first to report that sediment microbial communities exhibit significant beta diversity between gravel bars and the free-flowing reaches, implying that environmental heterogeneity due to in-channel modifications drive microbial community composition in this system.

Furthermore, we found a significant positive correlation between the functional diversity of the identified microbial taxa and beta diversity, suggesting that differences in the detected metabolic functions were closely related to dissimilarities in community composition. Likewise, closely associated variations in taxonomic and functional composition have been reported on eutrophic river-lake water and sediment bacterial communities [40], and for marine microbial communities on both temporal and spatial scales [16, 41]. Although the functional assignment based on microbial taxonomy employed in this study is putative, our results provide valuable insights into the potential microbial processes in the sediment that might be contributing to the biogeochemical processes carried out by the microbial communities in the Trinity River in comparison to its pristine tributaries and present its linear relationship with increasing community dissimilarity owing to the re-establishment of environmental heterogeneity due to gravel bar restoration.

*Proteobacteria*, *Bacteroidetes*, and *Acidobacteria*, which have been previously described as ubiquitous freshwater lineages using clone-based library approaches [42–44], were the highly abundant sediment microbial phyla on most samples, specifically in T-B. Functional annotation assigned most of the representatives of these phyla as chemoheterotrophs and aerobic chemoheterotrophs, hence, aerobic, chemoheterotrophy were the most abundant functional groups annotated for all samples, with T-B having relatively higher representations (Fig. 4). Notable genera classified under these functional groups were *Hyphomicrobia* and *Sphingomonas* under *Proteobacteria*, *Flavobacterium* (*Bacteroidetes*), and *Bryobacteria* (*Acidobacteria*). Substantial aerobic, chemoheterotrophic bacteria were numerically predominant components of the deep surface microflora of a southeast coastal plain [45]. Heterotrophic bacteria are often associated with decomposition and are responsible for the *in situ* pollution repair and degradation of organic matter [46–47]. Stegen et al. [48] reported that the ground and surface water mixing in the hyporheic zone primarily stimulates heterotrophic respiration, altering organic carbon composition, causes shifts from stochastic to deterministic ecological processes, and is associated with abundant microbial taxa with abilities to degrade a broad suite of organic compounds. The abundance of heterotrophic annotation of the taxa detected in T-B implies that these in-channel structures provide suitable areas for heterotrophic microorganisms to function, contributing to the net respiration in the river ecosystem. Accordingly, we identified *Flavobacterium*, *Sediminibacterium*, and members of the family *Rhodobacteraceae* as indicator taxa associated with T-B. *Flavobacterium* contains members capable of mineralizing various types of organic compounds [49], whereas members of the *Rhodobacteraceae* commonly found in freshwater environments [50] were detected in sediments where nitrate and iron reduction occurred [51]. Phototrophic *Rhodobacteraceae* were also known to perform denitrification function [52]. However, members of the genus *Sediminibacterium* from the family *Chitinophagaceae* which have been isolated from various sediment sources such as eutrophic [53] and freshwater reservoirs [54] are not yet well characterized in terms of their ecological role in aquatic systems.

Our results further revealed that most of the taxa identified in T-B had been annotated with aerobic nitrite oxidation and nitrification functions. Also, T-F reaches have notable nitrate and nitrogen respiration functions. These observations were supported by the dbRDA between the microbial communities and the environmental parameters presenting the negative relationship between NH_4_-N and NO_3_-N concentrations. NO_3_-N concentration increased in the gravel bar of the main channel, whereas NH_4_-N concentration was relatively higher in the free-flowing reaches of both the Trinity River and its tributaries. The microbes responsible for the aerobic transformation of nitrite to nitrate must have been the reason for the increase in NO_3_-N concentrations on the T-B reaches. Additionally, NO_3_-N is the first alternative electron acceptor utilized after oxygen in aquatic ecosystems [55], which implies that anaerobic metabolism was also induced in these reaches. In-stream geomorphic features, i.e., pools and riffles of urban restored and degraded sites, have been reported as strong sinks for nitrogen by maintaining anaerobic conditions and microbial biomass and activity [56]. NH_4_-N can result from dissimilatory NO_3_-N reduction by the anaerobic respiration of chemoheterotrophic microbes suggesting that these anaerobic processes were mostly promoted in T-F and R-F reaches. Also, the free-flowing reaches (both the T-F and R-F) identified *Betaproteobacteriales* (*Gammaproteobacteria*) as indicator taxa, which include metanotrophic and methylotrophic microorganisms associated with aerobic methane oxidation [57]. Accordingly, functional annotation revealed methanol oxidation, and methylotrophic functions were detected in the free-flowing communities.

Previous studies have shown that environmental factors are involved in shaping microbial community structure in aquatic environments [58–59]. Identifying the distribution patterns and the primary environmental drivers influencing microbial assemblage would provide a more comprehensive understanding of lotic ecosystem responses to changes in the environment [60–61]. As shown in the dbRDA analysis, pH and AFDM are the environmental predictors that significantly best explains the variability in microbial community composition (Fig. 3). Our findings on AFDM was consistent with Ock et al. [5] which reported that the restored gravel bars at the Trinity River have reduced suspended particulate organic matter concentration (i.e., μg of AFDM/ liter of water filtered) at base flow, with increased suspended particulate organic matter retention. Coarse particulate organic matter serves as the substrate for microbial activity promoting nutrient retention in streams [62], and functions as the primary carbon source for sediment and interstitial bacteria in the riparian zone of a river [63]. Sediments serve as an effective organic matter sink, and sediment microbial activity highly influences organic carbon biogeochemistry which could be greater for rivers with diverse channel morphology, including dynamic sediment structures favoring hydrodynamic exchange and residence time distribution [64] as provided by gravel bars [6, 9]. Additionally, the pH level significantly influenced the distribution of the sediment microbiome and was found to be significantly different between the Trinity River and tributary gravel bars, and free-flowing sites. This observation was consistent with previous studies in freshwater sediments reporting that variations in bacterial communities within wetland sediments were highly influenced by pH [22, 65]. Almost all of the Trinity River gravel bar points showed circumneutral pH levels except for one (i.e., B5DZ). Microbial diversity in terrestrial soil habitats strongly influenced by pH showed higher diversity with neutral pH than high or low ones [66].

On the other hand, although the Trinity River communities (by “River”) and the gravel bar communities (by “Reach”) were seen to have relatively similar trends of higher alpha diversity in comparison to the tributary and free-flowing samples, almost no significant difference among the 24 sediment microbial communities were observed except for the Simpson and Shannon indices by “River”. Our findings were consistent with previous studies that reported shifts in microbial community structure in urban streams, but no impacts on alpha diversity were observed through relative diversity indices [58, 66–67]. Also, some studies reported no difference in bacterial richness and a decline in diversity between rivers that have undergone restoration in comparison to degraded rivers [68]. The similar levels of with-in group diversity could be maintained across the urbanization gradient due to the introduction of novel bacterial taxa, which compensates for the loss of taxa sensitive to urban conditions [69].

## Conclusion

We utilized microbial communities as indicators of environmental changes to evaluate the progression of in-stream habitat restoration activities in the Trinity River, California, in particular, the restoration of gravel bars along the dam-impounded main channel. The compositions of the sediment microbial communities from the free-flowing sites of the Trinity River and its tributaries were relatively similar to one another, while the gravel bar communities from the main channel differed significantly with the tributaries. Construction or restoration of gravel bars contributed to the increased microbial biodiversity through the restoration of environmental heterogeneity at the river scale. This provides supporting evidence on the positive impact of habitat restoration in the Trinity River with the non-dam influenced, undisturbed tributaries as the basis of comparison. *Proteobacteria*, *Bacteroidetes*, and *Acidobacteria* were the highly abundant sediment microbial phyla on most sites, specifically in the Trinity River gravel bars sites. Moreover, chemoheterotrophy and aerobic chemoheterotrophy were the most prevalent microbial processes, with the Trinity River gravel bar reaches having relatively higher representations. The considerably large abundance of these heterotrophic taxa implies that gravel bars provide suitable areas for heterotrophic microorganisms with metabolic functions contributing to the net respiration in the riverine ecosystem. We report that pH and AFDM significantly best explain the variability in microbial community composition. It is also essential to note that the observed negative relationship between NH_4_-N and NO_3_-N concentrations were reflected in the functional annotation of Trinity River gravel bars’ aerobic nitrite oxidation and nitrification functions, and the Trinity River free-flowing reaches with nitrate and nitrogen respiration. Moreover, we report the significant positive correlation of taxonomic and functional diversity, implying that changes in sediment microbial community dissimilarity due to habitat heterogeneity also influenced the functional metabolic potential of the communities. Although functional assignment based on microbial taxonomy is putative, our results provide valuable insights into the potential microbial processes in the sediment that might be contributing to the biogeochemical processes carried out by the microbial communities in the Trinity River in comparison to its pristine tributaries.

## Materials and methods

### Study site and sample collection

The field survey was conducted in the Trinity River along a 60-km river length downstream of the Lewiston Dam in August 2017 (Fig. S1). Six gravel bars were sampled at the head (down-welling zone; DZ), and tail (up-welling zone; UZ), and two free-flowing segments were sampled at the up- (US) and downstream (DS) points along the Trinity River channel. The gravel bars were assessed correctly, and only those sites with distinct down-welling surface water and upwelling groundwater were sampled. To assess and compare dam-impacted gravel bars from the main channel against non-dam influenced areas, two gravel bars from tributaries, i.e., the Rush Creek and the Canyon Creek, were assessed. Also, two free-flowing segments from the tributary sites, as mentioned earlier, were sampled to provide a baseline comparison between the gravel bars and the non-in-gravel feature segments of the river. In total, twenty-four sampling points were sampled in this study.

Sediment samples were collected approximately 10 cm below the submerged surface of the gravel bars and the free-flowing areas using 50-mL sterile falcon tubes and immediately stored in 100% ethanol upon collection. Physical and chemical characteristics were measured from each of the sampling points. Dissolved oxygen (DO) was measured using a DO meter (Model OM-51, Horiba, Ltd., Kyoto, Japan). pH and electric conductivity (EC) was measured via the LAQUAtwin COND (Model Y071L, Horiba) and pH meters (Model Y017, Horiba). Water samples were collected following the procedures instructed by USEPA [70] and kept in an icebox in the field before sending it to PHYSIS Environmental Laboratories, Inc. (Anaheim, CA, USA) for water quality analysis. The chemical parameters assessed were ammonium nitrogen (NH_4_-N), nitrate nitrogen (NO_3_-N), ash-free dry mass (AFDM), and total suspended solids (TSS).

### Sequencing and data processing

Sediment samples for DNA extraction were homogenized and divided into triplicates of 200 mg each. The subsamples were then suspended in a buffer consisting of 10 mM EDTA, 50 mM Tris-HCl, 50 mM Na_2_HPO_4_·7H_2_O at pH 8.0 to remove PCR inhibitors [71–72]. Genomic DNA was extracted from the triplicate subsamples following the protocol of Zhou et al. [71] (as employed in 73) with some modifications, and the subsample extracts were combined before amplification. Library preparation was carried out through PCR using modified primer sequences targeting the V4 hypervariable region of the 16S SSU rRNA gene, i.e., 515F and 806R shown to yield optimal community analysis supported by the Earth Microbiome Project [74], with 12-base error-correcting Golay code on both forward and reverse primers. See supplementary text for the detailed amplicon library preparation (Additional file 1). A negative control sample was used to monitor any contamination from DNA extraction, and PCR to post-amplification library quantity and quality verification; however, no quantifiable amplicon was detected for further analysis. The normalized and pooled amplicon library was sequenced by the Advanced Research Support Center of Ehime University using the Illumina MiSeq platform with paired-end reads of 300-bp per read.

The raw sequence reads generated on the Illumina MiSeq platform were demultiplexed via the QIIME2 v2018.11 package [75]. Demultiplexed sequence data were quality screened, processed, and amplicon sequence variants (ASVs) were generated via the DADA2 v1.12 package in R [76], and classification was carried out via the SILVA ACT: Alignment, Classification and Tree Service (http://www.arb-silva.de/aligner) online server [77]. For the full read processing details, see the supplemental materials and methods section (Additional file 1). An interactive Krona chart was used to visualize the total relative abundances within the complex hierarchies of the microbial assignments [78].

Taxonomic heat trees were generated via *metacoder* [79] to pairwise-compare the four categorical groups based on “River × Reach”, and identify the taxa most strongly driving compositional differences based on ASV abundance. Moreover, indicator species analysis was performed to identify indicator taxa from the “River × Reach” category at the genus-level via the *multipatt* function implemented in the *indicspecies* package in R with 99,999 permutations [80].

The Functional Annotation of Prokaryotic Taxa (FAPROTAX 1.1) database was used to determine the potential microbial functions of the taxonomically assigned dataset [81]. Detected taxa were assigned to one or more established metabolic or ecologically relevant functions (e.g., nitrate respiration, methanogenesis, fermentation, or plant pathogenesis) based on available literature on cultured representatives within the taxon that has been reported to exhibit that function.

The raw sequence data were deposited into the National Center for Biotechnology Information (NCBI) Sequence Read Archive (SRA) and is available under BioProject accession code PRJNA559761.

### Statistical analysis

Homogeneity of multivariate dispersion (PERMDISP2) [82] implemented in the *betadisper* function of the *vegan* 2.5-5 package [83] under the R program (http://cran.r-project.org/) was used to assess if differences in environmental heterogeneity existed among the samples based on different categorical factors: (i) “River”, (ii) “Reach” (iii) “Point”, and their combinations (iv) “River × Reach”, and (v) “River × Reach × Point”. The resulting ANOVA *F*-statistic was used to compare among-group differences in distances between observations and group medians. Alpha diversity (within-sample diversity) and evenness estimates, i.e., Chao1, Fisher’s alpha, Shannon, and Simpson’s index, were calculated using QIIME. Before the analysis, the input table consisting of the ASVs with the accepted taxonomic assignment was rarefied to the sampling point with the lowest read abundance. The average values of each calculated alpha diversity metrics among different categories were tested using Student’s t-test for independent means or one-way analysis of variance (ANOVA) to facilitate comparison between the diversity estimates of the samples based on different categorical factors.

Read abundances were rarefied via the *rrarefy* function to the sampling point with the lowest abundance and log-transformed (“log1p” - natural logarithm of the value plus one) before the analyses. The differences in community structure (beta diversity) were assessed by conducting non-parametric analyses. The differences in community structure between groups based on the different categorical factors were tested using the analysis of similarity (ANOSIM) [84] via the *anosim* function. ANOSIM was used to assess if the similarity among groups sample is greater than within-group samples. A 3-way permutational multivariate analysis of variance using distance matrices (PERMANOVA) [85] (with 9,999 permutations, Bray-Curtis) implemented in the *adonis2* function was performed among the categorical factors, i.e., “River”, “Reach” and “Point” to test the statistical significance of the between-groups distances. Homogeneity of multivariate dispersion was also verified for each categorical factor (and combinations) with the distance-based test (PERMDISP2) [82] implemented in the *betadisper* function visualized with Principal Coordinates Analysis (PCoA) plots. Also, NMDS ordination based on Bray-Curtis distance was performed at the genus-level dataset for the 24 sampling points to show the dissimilarity (beta diversity) between the microbial communities among the samples.

Distance-based redundancy analysis (dbRDA) was performed to determine the environmental variables responsible for explaining community composition variation amongst sampling points with significance. The dbRDA plots were generated via the *vegan* package in R with the *cca* function. Before the analysis, environmental variables were evaluated for multi-collinearity via variance inflation factors (VIF) for constraining parameters (VIF <5) using the *usdm* package [86]. The significance of the correlation of each environmental variable with the microbial communities was determined by the permutation test and bootstrap with 1000 iterations.

Moreover, Mantel test was performed to assess how beta diversity (based on Bray-Curtis and Jaccard distance calculated from the dataset at the genus-level) correlated with functional dissimilarity (based on Jaccard distance for presence/absence estimation from the functional annotation data) via the *mantel* function with 9,999 permutations. Procrustes tests were also employed as an alternative to Mantel tests to compare the two matrices via the *protest* function with 9,999 permutations. Ordination and heatmap generation were carried out on the R software (version 3.6.0) under the *vegan* 2.5-5 [83] and *pheatmap* [84] packages.

## Supplementary information

**Additional file 1:** Krona chart of identified sediment microbial taxa. Browsable Krona chart, Additional_File1.html. (HTML 285 kb)

**Additional file 2: Fig. S1** Sampling location. **a** The Trinity River and its tributaries impounded by the Trinity Dam and the Lewiston Dam in California, USA. **b** Gravel bars were sampled at the head (down-welling zone; DZ), and tail (up-welling zone; UZ) points. **c** Free-flowing sites were assessed, and samples were collected on up-(US) and downstream (DS) points. **d** In total, eight gravel bars and four free-flowing sites were assessed in the study; gravel bars B1 and B4 drone photos shown as examples. Parts of the map was generated from Google Earth Pro (version 7.3.2.5776; https://www.google.com/earth/). **Fig. S2** The relative abundance of the microbial community composition of the 24 sampling points at the genus-level (showing taxa with >1% read abundance). **Fig. S3** The relative abundance of the microbial community composition of the 24 sampling points at the phylum-level. **Fig. S4** Alpha diversity estimates (mean value) for each categorical factors. **Fig. S5** Microbial community composition at the genus-level of all 24 sampling points indicated by a non-metric multidimensional scaling plot (NMDS). **Fig. S6** Box plots of distances to group centroids from PERMANOVA (A-E) and PERMDISP2 (F-G). **Fig. S7** Pairwise taxonomic comparisons (ASVs with relative read abundance >0.10) between the “Reach” categories. **Fig. S8** The relative abundance of the predicted functional groups of the 24 sampling points. **Table S1** Physico-chemical properties. **Table S2** PERMDISP2 analysis to test the effect of the categorical factors on the physicochemical parameters. **Table S3** Read processing and amplicon sequence variants (ASVs) generation. **Table S4** Richness and diversity estimates of the 24 sampling points. **Table S5** Multivariate pattern analysis for indicator taxa. (DOCX 5803 kb)

## Abbreviations

TRRP: Trinity River Restoration Program;
T: Trinity River;
R: Reference tributaries;
B: Gravel bar;
F: Free-flowing segment;
UZ: Up-welling zone of gravel bar;
DZ: Down-welling of gravel bar;
US: upstream of free-flowing segment;
DS: downstream of free-flowing segment;
DO: Dissolved oxygen;
NH_4_-N: Ammonium nitrogen;
NO_3_-N: Nitrate nitrogen;
AFDM: Ash-free dry mass;
TSS: Total suspended solids;
ASVs: Amplicon sequence variants;
FAPROTAX: Functional annotation of prokaryotic taxa;
VIF: Variance inflation factors;
NMDS: Nonmetric multidimensional scaling;
dbRDA: Distance-based redundancy analysis;
PCoA: Principal coordinates analysis;
ANOSIM: analysis of similarity;
PERMANOVA: Permutational multivariate analysis of variance.

## Acknowledgments

The authors are grateful to the members of the Disaster Prevention Research Institute (DPRI), Kyoto University, and Dr. David Gaeuman of the Trinity River Restoration Program for their assistance during the field survey. We thank Dr. Naohito Tokunaga of the Division of Analytical Bio-Medicine for his assistance on performing high-throughput sequencing on the Illumina MiSeq platform of the Advanced Research Support Center (ADRES), Ehime University.

## Author’s contributions

KW and JMS conceptualized this study. JS, BL, YT, and TS conducted fieldwork. JS performed laboratory and molecular data analysis. Data interpretation was performed by JS and KW. JS wrote the main manuscript text. KW did critical revisions of the manuscript. All authors reviewed and approved the final version of the manuscript.

## Availability of data and materials

The raw Illumina sequence data were deposited into the National Center for Biotechnology Information (NCBI) Sequence Read Archive (SRA) under BioProject with the accession number PRJNA559761.

## Ethics approval and consent to participate

Not applicable.

## Consent for publication

Not applicable.

## References

1. Yan Q, Bi Y, Deng Y, He Z, Wu L, Van Nostrand JD, Yu Y, et al. Impacts of the Three Gorges Dam on microbial structure and potential function. Sci Rep. 2015;5:8605.

2. Lansac-Tôha FM, Heino J, Quirino BA, Moresco GA, Peláez O, Meira BR, Velho LFM, et al. Differently dispersing organism groups show contrasting beta diversity patterns in a dammed subtropical river basin. Sci Total Environ. 2019;691:1271–1281.

3. Chaparro G, O’Farrell I, Hein T. Multi-scale analysis of functional plankton diversity in floodplain wetlands: Effects of river regulation. Sci Total Environ. 2019;667:338–347.

4. Kondolf GM, Anderson S, Lave R, Pagano L, Merenlender A, Bernhardt ES. Two decades of river restoration in California: What can we learn? Restor Ecol. 2007;15(3):516–523.

5. Ock G, Gaeuman D, McSloy J, Kondolf GM. Ecological functions of restored gravel bars, the Trinity River, California. Ecol Eng. 2015;83:49–60.

6. Beechie TJ., Sear DA, Olden JD, Pess GR, Buffington JM, Moir H, Pollock M, et al. Process-based principles for restoring river ecosystems. BioScience. 2010;60(3):209–222.

7. Boano F, Harvey JW, Marion A, Packman AI, Revelli R, Ridolfi L, Wörman A. Hyporheic flow and transport processes: Mechanisms, models, and biogeochemical implications. Rev Geophys. 2014;52(4):603–679.

8. Hester ET, Doyle MW. In-stream geomorphic structures as drivers of hyporheic exchange. Water Resour Res. 2008;44:3.

9. Sackett JD, Shope CL, Bruckner JC, Wallace J, Cooper CA, Moser DP. Microbial Community Structure and Metabolic Potential of the Hyporheic Zone of a Large Mid-Stream Channel Bar. Geomicrobiol J. 2019;36(9):765–776.

10. Mendoza-Lera C, Datry T. Relating hydraulic conductivity and hyporheic zone biogeochemical processing to conserve and restore river ecosystem services. Sci Total Environ. 2017;579:1815–1821.

11. Leibold MA, Holyoak M, Mouquet N, Amarasekare P, Chase JM, Hoopes MF, Loreau M, et al. The metacommunity concept: a framework for multi-scale community ecology. Ecol Lett. 2004;7(7):601–613.

12. Soininen J, Heino J, Wang J. A meta-analysis of nestedness and turnover components of beta diversity across organisms and ecosystems. Glob Ecol Biogeogr. 2018;27(1):96–109.

13. Heino J, Grönroos M, Ilmonen J, Karhu T, Niva M, Paasivirta L. Environmental heterogeneity and β diversity of stream macroinvertebrate communities at intermediate spatial scales. Freshw Sci. 2012;32(1):142–154.

14. Lowell JL, Gordon N, Engstrom D, Stanford JA, Holben W E, Gannon JE. Habitat heterogeneity and associated microbial community structure in a small-scale floodplain hyporheic flow path. Microb Ecol. 2009;58(3):611–620.

15. Ramond P, Sourisseau M, Simon N, Romac S, Schmitt S, Rigaut-Jalabert F, Siano R, et al. Coupling between taxonomic and functional diversity in protistan coastal communities. Environ Microbiol. 2019;21(2):730–749.

16. Galand PE, Pereira O, Hochart C, Auguet JC, Debroas D. A strong link between marine microbial community composition and function challenges the idea of functional redundancy. ISME J. 2018;12(10):2470.

17. Fasching C, Akotoye C, Bižić M, Fonvielle J, Ionescu D, Mathavarajah S, Xenopoulos MA, et al. Linking stream microbial community functional genes to dissolved organic matter and inorganic nutrients. Limnol Oceanogr. 2020;doi.org/10.1002/lno.11356

18. Staley C, Gould TJ, Wang P, Phillips J, Cotner JB, Sadowsky MJ. Species sorting and seasonal dynamics primarily shape bacterial communities in the Upper Mississippi River. Sci Total Environ. 2015;505:435–445.

19. Friberg N. Impacts and indicators of change in lotic ecosystems. WIRES Water. 2014;1(6):513–531.

20. Tremblay L, Benner R. Organic matter diagenesis and bacterial contributions to detrital carbon and nitrogen in the Amazon River system. Limnol Oceanogr. 2009;54(3):681–691.

21. Boulton AJ, Findlay S, Marmonier P, Stanley EH, Valett HM. The functional significance of the hyporheic zone in streams and rivers. Annu Rev Ecol Syst. 1998;29(1):59–81.

22. Liu S, Ren H, Shen L, Lou L, Tian G, Zheng P, Hu B. pH levels drive bacterial community structure in sediments of the Qiantang River as determined by 454 pyrosequencing. Front Microbiol. 2015;6:285.

23. Boulton AJ. Hyporheic rehabilitation in rivers: restoring vertical connectivity. Freshw Biol. 2007;52(4):632–650.

24. Olsen DA, Townsend CR. Hyporheic community composition in a gravel-bed stream: influence of vertical hydrological exchange, sediment structure and physicochemistry. Freshw Biol. 2003;48(8):1363–1378.

25. Nogaro G, Datry T, Mermillod-Blondin FL, Descloux S, Montuelle B. Influence of streambed sediment clogging on microbial processes in the hyporheic zone. Freshw Biol. 2010;55(6):1288–1302.

26. Chen J, Wang P, Wang C, Wang X, Miao L, Liu S, Yuan Q. Bacterial communities in riparian sediments: a large-scale longitudinal distribution pattern and response to dam construction. Front Microbiol. 2018;9:999.

27. Shepherd RG. Correlations of permeability and grain size. Groundwater. 1989;27(5):633–638.

28. Gayraud S, Philippe M. Influence of bed-sediment features on the interstitial habitat available for macroinvertebrates in 15 French streams. Int Rev Hydrobiol. 2003;88(1):77–93.

29. Nogaro G, Datry T, Mermillod-Blondin F, Foulquier A, Montuelle B. Influence of hyporheic zone characteristics on the structure and activity of microbial assemblages. Freshw Biol. 2013;58(12):2567–2583.

30. Levi PS, Starnawski P, Poulsen B, Baattrup-Pedersen A, Schramm A, Riis T. Microbial community diversity and composition varies with habitat characteristics and biofilm function in macrophyte-rich streams. Oikos. 2017;126(3):398–409.

31. Lemke MJ, Paver SF, Dungey KE, Velho LFM, Kent AD, Rodrigues LC, Randle MR, et al. Diversity and succession of pelagic microorganism communities in a newly restored Illinois River floodplain lake. Hydrobiologia. 2017;804(1):35–58.

32. Gaeuman D. High-flow gravel injection for constructing designed in-channel features. River Res Appl. 2014;30(6):685–706.

33. Gaeuman D, Stewart R, Schmandt B, Pryor C. Geomorphic response to gravel augmentation and high-flow dam release in the Trinity River, California. Earth Surf Process Landf. 2017;42(15):2523–2540.

34. Trauth N, Schmidt C, Vieweg M, Oswald SE, Fleckenstein JH. Hydraulic controls of in-stream gravel bar hyporheic exchange and reactions. Water Resour Res. 2015;51(4):2243–2263.

35. Claret C, Marmonier P, Boissier JM, Fontvieille D, Blanc P. Nutrient transfer between parafluvial interstitial water and river water: influence of gravel bar heterogeneity. Freshw Biol. 1997;37(3):657–670.

36. Miller SW. Budy P, Schmidt JC. Quantifying macroinvertebrate responses to in-stream habitat restoration: applications of meta-analysis to river restoration. Restor Ecol. 2010;18(1):8–19.

37. Harrison SSC, Pretty JL, Shepherd D, Hildrew AG, Smith C, Hey RD. The effect of instream rehabilitation structures on macroinvertebrates in lowland rivers. J Appl Ecol. 2004;41(6):1140–1154.

38. Singer G, Besemer K, Schmitt-Kopplin P, Hödl I, Battin TJ. Physical heterogeneity increases biofilm resource use and its molecular diversity in stream mesocosms. PLoS One. 2010;5(4):e9988.

39. Besemer K, Singer G, Hödl I, Battin TJ. Bacterial community composition of stream biofilms in spatially variable-flow environments. Appl Environ Microbiol. 2009;75(22):7189–7195.

40. Ren Z, Qu X, Peng W, Yu Y, Zhang M. Functional properties of bacterial communities in water and sediment of the eutrophic river-lake system of Poyang Lake, China. PeerJ. 2019;7:e7318.

41. Galand PE, Salter I, Kalenitchenko D. Ecosystem productivity is associated with bacterial phylogenetic distance in surface marine waters. Mol Ecol. 2015;24(23):5785–5795.

42. Tamaki H, Sekiguchi Y, Hanada S, Nakamura K, Nomura N, Matsumura M, Kamagata Y. Comparative analysis of bacterial diversity in freshwater sediment of a shallow eutrophic lake by molecular and improved cultivation-based techniques. Appl Environ Microbiol. 2005;71(4):2162–2169.

43. Jones RT, Robeson MS, Lauber CL, Hamady M, Knight R, Fierer N. A comprehensive survey of soil acidobacterial diversity using pyrosequencing and clone library analyses. ISME J. 2009;3(4):442.

44. Staley C, Unno T, Gould TJ, Jarvis B, Phillips J, Cotner JB, Sadowsky MJ. Application of Illumina next-generation sequencing to characterize the bacterial community of the Upper Mississippi River. Journal of Applied Microbiology. 2013;115(5):1147–1158.

45. Balkwill DL. Numbers, diversity, and morphological characteristics of aerobic, chemoheterotrophic bacteria in deep subsurface sediments from a site in South Carolina. Geomicrobiol J. 1989;7(1-2):33–52.

46. Freixa A, Ejarque E, Crognale S, Amalfitano S, Fazi S, Butturini A, Romaní AM. Sediment microbial communities rely on different dissolved organic matter sources along a Mediterranean river continuum. Limnol Oceanogr. 2016;61(4):1389–1405.

47. Wei Z, Liu Y, Feng K, Li S, Wang S, Jin D, Deng Y, et al. The divergence between fungal and bacterial communities in seasonal and spatial variations of wastewater treatment plants. Sci Total Environ. 2018;628:969–978.

48. Stegen JC, Fredrickson JK, Wilkins MJ, Konopka AE, Nelson WC, Arntzen EV, Kennedy DW, et al. Groundwater–surface water mixing shifts ecological assembly processes and stimulates organic carbon turnover. Nat Commun. 2016;7:11237.

49. McBride MJ. The family Flavobacteriaceae. In: Rosenberg, E., et al. (Eds.), The Prokayotes. Springer Berlin Heidelberg, Berlin, Heidelberg; 2014. p. 439–512.

50. Pujalte MJ, Lucena T, Ruvira MA, Ruiz Arahal D, Carmen Macian M. The family Rhodobacteraceae. In: Rosenberg, E., et al. (Eds.), The Prokayotes. Springer Berlin Heidelberg, Berlin, Heidelberg; 2014. p. 439–512.

51. Reyes C, Dellwig O, Dähnke K, Gehre M, Noriega-Ortega BE, Böttcher ME, Friedrich MW, et al. Bacterial communities potentially involved in iron-cycling in Baltic Sea and North Sea sediments revealed by pyrosequencing. FEMS Microbiol Ecol. 2016;92(4):fiw054.

52. Shapleigh JP. Denitrifying prokaryotes: Alphaproteobacteria. E. Rosenberg, E.F DeLong, S. Lory, E. Stackebrandt, F. Thompson, editors. In: The Prokaryotes-Prokaryotic Physiology and Biochemistry. Berlin, Springer-Verlag; 2013. p. 405–425.

53. Qu JH, Yuan HL. *Sediminibacterium salmoneum* gen. nov., sp. nov., a member of the phylum Bacteroidetes isolated from sediment of a eutrophic reservoir. Int J Syst Evol Microbiol. 2008;58(9):2191–2194.

54. Kang H, Kim H, Lee BI, Joung Y, Joh K. *Sediminibacterium goheungense* sp. nov., isolated from a freshwater reservoir. Int J Syst Evol Microbiol. 2014;64(4):1328–1333.

55. Steinberg NA, Blum JS, Hochstein L, Oremland RS. Nitrate is a preferred electron acceptor for growth of freshwater selenate-respiring bacteria. Appl Environ Microbiol. 1992;58(1):426–428.

56. Harrison MD, Groffman PM, Mayer PM, Kaushal SS. Microbial biomass and activity in geomorphic features in forested and urban restored and degraded streams. Ecol Eng. 2012;38(1):1–10.

57. Taubert M, Grob C, Crombie A, Howat AM, Burns OJ, Weber M, von Bergen M, et al. Communal metabolism by *Methylococcaceae* and *Methylophilaceae* is driving rapid aerobic methane oxidation in sediments of a shallow seep near Elba, Italy. Environ Microbiol. 2019;21(10):3780–3795.

58. Zeglin LH. Stream microbial diversity in response to environmental changes: review and synthesis of existing research. Front Microbiol. 2015;6:454.

59. Yao L, Chen C, Liu G, Li F, Liu W. Environmental factors, but not abundance and diversity of nitrifying microorganisms, explain sediment nitrification rates in Yangtze lakes. RSC Adv. 2018;8(4):1875–1883.

60. Lau KE, Washington VJ, Fan V, Neale MW, Lear G, Curran J, Lewis GD. A novel bacterial community index to assess stream ecological health. Freshw Biol. 2015;60(10):1988–2002.

61. Liao H, Yu K, Duan Y, Ning Z, Li B, He L, Liu C. Profiling microbial communities in a watershed undergoing intensive anthropogenic activities. Sci Total Environ. 2019;647:1137–1147.

62. Aldridge KT, Brookes JD, Ganf GG. Rehabilitation of stream ecosystem functions through the reintroduction of coarse particulate organic matter. Restor Ecol. 2009;17(1):97–106.

63. Brugger A, Wett B, Kolar I, Reitner B, Herndl GJ. Immobilization and bacterial utilization of dissolved organic carbon entering the riparian zone of the alpine Enns River, Austria. Aquat Microb Ecol. 2001;24(2):129–142.

64. Fischer H, Sachse A, Steinberg CE, Pusch M. Differential retention and utilization of dissolved organic carbon by bacteria in river sediments. Limnol Oceanogr. 2002;47(6):1702–1711.

65. Ligi T, Oopkaup K, Truu M, Preem JK, Nõlvak H, Mitsch WJ, Truu J, et al. Characterization of bacterial communities in soil and sediment of a created riverine wetland complex using high-throughput 16S rRNA amplicon sequencing. Ecol Eng. 2014;72:56–66.

66. Lauber CL, Hamady M, Knight R, Fierer N. Pyrosequencing-based assessment of soil pH as a predictor of soil bacterial community structure at the continental scale. Appl Environ Microbiol. 2009;75(15):5111–5120.

67. Roberto AA, Van Gray JB, Leff LG. Sediment bacteria in an urban stream: spatiotemporal patterns in community composition. Water Res. 2018;134:353–369.

68. Simonin M, Voss KA, Hassett BA, Rocca JD, Wang SY, Bier RL, Bernhardt ES, et al. In search of microbial indicator taxa: shifts in stream bacterial communities along an urbanization gradient. Environ Microbiol. 2019;doi:10.1111/1462-2920.14694.

69. Lin Q, Sekar R, Marrs R, Zhang Y. Effect of River Ecological Restoration on Biofilm Microbial Community Composition. Water. 2019;11(6):1244.

70. Hosen JD, Febria CM, Crump BC, Palmer MA. Watershed urbanization linked to differences in stream bacterial community composition. Front Microbiol. 2017;8:1452.

71. USEPA. Integrated Risk Information System (IRIS). 2014; http://www.epa.gov/iris/. Accessed 01 August 2017.

72. Zhou J, Bruns MA, Tiedje JM. DNA recovery from soils of diverse composition. Appl Environ Microbiol. 1996;62(2):316–322.

73. Poulain AJ, Aris-Brosou S, Blais JM, Brazeau M, Keller WB, Paterson AM. Microbial DNA records historical delivery of anthropogenic mercury. ISME J. 2015;9(12):2541.

74. Solomon S, Kachiprath B, Jayanath G, Sajeevan TP, Singh IB, Philip R. High-quality metagenomic DNA from marine sediment samples for genomic studies through a preprocessing approach. 3 Biotech. 2016;6(2):160.

75. Caporaso JG, Lauber CL, Walters WA, Berg-Lyons D, Huntley J, Fierer N, Gormley N, et al. Ultra-high-throughput microbial community analysis on the Illumina HiSeq and MiSeq platforms. ISME J. 2012;6(8):1621.

76. Bolyen E, Rideout JR, Dillon MR, Bokulich NA, Abnet CC, Al-Ghalith GA, Bai Y, et al. Reproducible, interactive, scalable and extensible microbiome data science using QIIME 2. Nat Biotechnol. 2019; doi.org/10.1038/s41587-019-0209-9.

77. Callahan BJ, McMurdie PJ, Rosen MJ, Han AW, Johnson AJA, Holmes SP. DADA2: high-resolution sample inference from Illumina amplicon data. Nat Methods. 2016;13(7):581.

78. Pruesse E, Peplies J, Glöckner FO. SINA: accurate high-throughput multiple sequence alignment of ribosomal RNA genes. Bioinformatics. 2012;28(14):1823–1829.

79. Ondov BD, Bergman NH, Phillippy AM. Interactive metagenomic visualization in a Web browser. BMC bioinformatics. 2011;12(1):385.

80. Foster ZS, Sharpton TJ, Grünwald NJ. Metacoder: An R package for visualization and manipulation of community taxonomic diversity data. PLoS Comput. 2017;13(2):e1005404.

81. De Caceres M, Jansen F, De Caceres MM. Package ‘indicspecies’. Relationship between species and groups of sites. R package. 2016;1:6.

82. Louca S, Parfrey LW, Doebeli M. Decoupling function and taxonomy in the global ocean microbiome. Sci. 2016;353(6305):1272–1277.

83. Anderson MJ. Distance-based tests for homogeneity of multivariate dispersions. Biometrics. 2006;62(1):245–253.

84. Oksanen J, Blanchet FG, Kindt R, Legendre P, Minchin PR, O’hara RB, Oksanen MJ, et al. Package ‘vegan’. Community ecology package. 2013;2(9):1–295.

85. Clarke KR. Non-parametric multivariate analyses of changes in community structure. Aust J Ecol. 1993;18(1):117–143.

86. Kolde R, Kolde MR. Package ‘pheatmap’. R Package. 2015;1:7.

87. Anderson MJ. A new method for non-parametric multivariate analysis of variance. Austral Ecol. 2001;26(1):32–46.

88. Naimi B. usdm: Uncertainty analysis for species distribution models. R Package. 2015;1:1–12.

