## Additional File 1 for "Profiling the microbial community structure and functional diversity of a dam-regulated river undergoing gravel bar restoration"

Javascript must be enabled to view this page.

magnitude

 905362

 537214

 234317

 207926

 113255

 103026

 9929

 271

 29

 58154

 56427

 1727

 14558

 14558

 11745

 3881

 6725

 862

 180

 84

 13

 3331

 3132

 117

 82

 1611

 1611

 2969

 2969

 1623

 889

 698

 36

 634

 634

 46

 46

 4879

 4879

 4879

 5686

 4977

 3610

 1233

 134

 709

 709

 1693

 1693

 1586

 66

 41

 2829

 2829

 2829

 1152

 1152

 1152

 2167

 1356

 1356

 765

 765

 46

 46

 853

 853

 853

 2034

 2034

 1382

 486

 166

 1252

 1252

 1252

 1323

 1323

 1323

 548

 548

 548

 617

 617

 617

 411

 246

 229

 17

 141

 125

 16

 24

 24

 222

 222

 222

 118

 118

 118

 325

 325

 325

 103

 103

 103

 92

 92

 92

 42

 42

 42

 27

 27

 27

 18

 18

 18

 263261

 83163

 83163

 55190

 20257

 2001

 2278

 3275

 137

 25

 78408

 32093

 22292

 7964

 1837

 21032

 21032

 2763

 2632

 117

 14

 2892

 2090

 780

 22

 6745

 5098

 1428

 171

 48

 7336

 6809

 401

 126

 2868

 2868

 1108

 1108

 1528

 1165

 321

 42

 27

 27

 16

 16

 64213

 64213

 60250

 3223

 558

 140

 42

 21916

 17869

 7124

 7554

 2977

 140

 74

 3158

 2818

 255

 39

 46

 889

 889

 2385

 2058

 2058

 149

 149

 159

 159

 19

 19

 6564

 6564

 6564

 2273

 1976

 1976

 297

 297

 1554

 1554

 935

 619

 537

 537

 537

 1046

 1046

 1046

 283

 283

 283

 148

 148

 148

 464

 464

 243

 152

 42

 27

 91

 60

 60

 22

 22

 9

 9

 32

 32

 32

 112

 35

 35

 77

 77

 35

 35

 21

 14

 19

 19

 19

 14

 14

 14

 4

 4

 4

 38869

 11262

 11262

 11262

 4684

 4684

 4684

 3120

 2759

 2627

 132

 361

 303

 58

 12109

 2965

 2965

 2092

 2092

 933

 933

 823

 823

 614

 509

 105

 1140

 1140

 2227

 1339

 431

 140

 317

 339

 339

 509

 509

 347

 347

 73

 73

 47

 47

 4487

 4487

 4487

 1377

 392

 392

 985

 985

 721

 570

 415

 136

 19

 129

 129

 22

 22

 300

 300

 300

 229

 229

 229

 329

 329

 329

 128

 114

 105

 9

 14

 14

 72

 72

 72

 34

 34

 34

 9

 9

 9

 8

 8

 8

 767

 767

 767

 767

 37987

 37250

 37250

 37250

 37200

 50

 507

 507

 507

 507

 230

 230

 230

 230

 164031

 163453

 85818

 31626

 31626

 53524

 53524

 459

 395

 64

 209

 209

 21242

 10294

 6471

 2535

 1288

 8708

 8250

 458

 1245

 1203

 42

 962

 962

 33

 33

 43142

 12718

 7156

 4799

 507

 256

 28806

 22830

 1850

 4117

 9

 648

 648

 970

 970

 13130

 6919

 6919

 2258

 2258

 2930

 2930

 417

 333

 84

 486

 486

 120

 120

 71

 71

 71

 50

 35

 35

 12

 12

 3

 3

 462

 361

 361

 361

 101

 101

 101

 116

 116

 116

 116

 34210

 34210

 14634

 10760

 10760

 3874

 3874

 11594

 11594

 11594

 6399

 6399

 6399

 1227

 1227

 1227

 199

 199

 199

 157

 157

 157

 32842

 11191

 11191

 11191

 11191

 7194

 7194

 7194

 7194

 3732

 3375

 3375

 3148

 227

 124

 124

 124

 109

 109

 109

 70

 70

 70

 54

 54

 54

 10276

 5859

 2748

 2519

 229

 3028

 3028

 83

 67

 16

 3378

 3378

 2751

 627

 418

 418

 324

 94

 577

 577

 465

 100

 6

 6

 44

 44

 44

 414

 414

 414

 414

 35

 35

 35

 35

 65097

 42312

 8506

 8506

 8506

 23173

 23173

 20818

 2355

 10633

 10633

 10633

 4852

 4852

 4852

 4852

 2514

 2366

 2366

 2280

 81

 5

 148

 148

 148

 1515

 1515

 1515

 1515

 12046

 12046

 12046

 12046

 1592

 1592

 1592

 1592

 162

 162

 162

 162

 67

 67

 67

 67

 30

 30

 30

 30

 7

 7

 7

 7

 7024

 7024

 5166

 5166

 5166

 599

 599

 599

 826

 826

 826

 128

 128

 128

 90

 90

 90

 112

 112

 112

 103

 87

 87

 8

 8

 6

 6

 2

 2

 3612

 2929

 2929

 2929

 2929

 497

 497

 497

 497

 186

 186

 186

 186

 3481

 2791

 2507

 2391

 1800

 591

 116

 116

 225

 225

 187

 38

 59

 59

 59

 690

 690

 690

 690

 1538

 1538

 1538

 1538

 1538

 8453

 8106

 8106

 8106

 6986

 1067

 53

 320

 320

 320

 320

 27

 27

 27

 27

 1087

 789

 789

 789

 789

 298

 298

 298

 298

 1051

 1025

 1025

 1025

 1025

 26

 26

 26

 26

 1326

 472

 472

 472

 472

 521

 443

 443

 443

 78

 78

 78

 333

 170

 170

 170

 130

 130

 130

 33

 33

 33

 423

 423

 423

 423

 423

 884

 884

 884

 884

 884

 1044

 1044

 1044

 1044

 1044

 948

 948

 948

 948

 948

 947

 947

 947

 947

 947

 341

 341

 341

 337

 271

 40

 26

 4

 4

 1076

 960

 267

 267

 267

 348

 348

 348

 174

 130

 130

 44

 44

 149

 73

 73

 76

 76

 22

 22

 22

 53

 53

 53

 53

 44

 44

 21

 21

 13

 13

 10

 10

 19

 19

 19

 19

 167

 167

 167

 167

 167

 198

 198

 198

 198

 198

 134

 134

 134

 134

 134

 195

 181

 181

 126

 126

 55

 55

 14

 14

 10

 10

 4

 4

 50

 50

 50

 50

 50

 2

 2

 2

 2

 2
