## Additional File 2 for "Profiling the microbial community structure and functional diversity of a dam-regulated river undergoing gravel bar restoration"

**Research Article**


**Authors and Affiliations**

Joeselle M. Serrana[1,2,3], Bin Li[1], Tetsuya Sumi[4], Yasuhiro Takemon[4], and Kozo Watanabe[1,2,3] *

[1]Department of Civil and Environmental Engineering, Ehime University, Bunkyo-cho 3, Matsuyama, Ehime 7908577, Japan

[2]Center for Marine Environmental Studies (CMES), Ehime University, Bunkyo-cho 2-5, Matsuyama, Ehime 7908577, Japan

[3]Biological Control Research Unit, Center for Natural Sciences and Environmental Research, De La Salle University, 2401 Taft Avenue, Manila 1004, Philippines

[4]Disaster Prevention Research Institute, Kyoto University, Gokasho, Uji, Kyoto 6110011, Japan

*****corresponding author

Prof. Kozo Watanabe, PhD

Phone & Fax Number: +81 (0) 89 927 9847

### Supplementary Text

### Materials and Methods

#### Site description

The Trinity River is a large gravel-bed river impounded by the Trinity Dam (164 m a.b.l. and 3020 million m3 storage) and the smaller Lewiston Dam (28 m a.b.l. and 18 million m3 storage) in northern California, USA under current dam operating guidelines with a mean annual flood of approximately 180 m3/s (Gaeuman et al., 2017). The Trinity River Restoration Program (TRRP), a multi-agency partnership, manages and implements these releases alongside gravel augmentations and mechanical rehabilitations downstream of the Lewiston Dam with the aims of restoring salmonid habitat and dynamic channel processes in the river (USDOI, 2000 as cited in Ock et al., 2015). In particular, TRRP conducts coarse-sediment augmentation to recreate or rehabilitate instream gravel features, i.e. gravel bars either via fluvial deposition of locally added sediments or by mechanical construction of gravel islands and bars.

#### PCR amplification and sequencing

Amplicon libraries were prepared using PCR with high-fidelity Phusion polymerase (Thermo Fisher Scientific Inc.) in a T100 Thermal Cycler (Bio-Rad Laboratories, USA). The 25 µl PCR reaction mixture consisted of 5 µl of 5X Phusion GC Buffer, 1.25 µl each of the forward and reverse primers (10 µM), two µl dNTPs (2.5 mM), 0.75 µl DMSO, 0.25 µl Phusion Polymerase (1 U) and one µl of template DNA. The PCR condition followed was initial denaturation at 98°C for 3 min, 25 cycles of denaturation at 98°C for 15 s, annealing at 50°C for 30 s and extension at 72°C for 30 s, followed by a final extension period at 72°C for 7 min. Post-amplification, library-quality control was performed by checking the library size distribution via the High-Sensitivity DNA chip (Agilent Bioanalyzer), and the libraries were purified/size selected using SPRI beads (AmpureXP, Beckman Coulter Genomics). Amplicon size was ~400 bp. The purified amplicon libraries were then quantified via the Kapa qPCR Illumina Quantification kit (Kapa Biosystems, USA), normalized, and equimolar amounts of the libraries from each sampling point were pooled. The 4 nM pooled library was sequenced by the Advanced Research Support Center (ADRES) of Ehime University using the Illumina MiSeq platform with paired-end reads of 300-bp per read.

Read processing and taxonomic assignment

Based on the read error profiles, the reverse reads have poor read quality. Low read abundance with acceptable overlaps between the reads can be accounted after quality filtering; therefore, only the forward reads were used in the subsequent analysis. Primer contaminants were excluded, and the reads were filtered based on quality, truncated at 100-bp, and identified sequence variants likely to be derived from sequencing error. Amplicon Sequence Variants (ASVs) (analogous to 100% sequence similarity in OTUs) were inferred from the sequence data, subsequently removing chimeric ASVs. The ASV sequences were then extracted and aligned to multiple databases using the SILVA ACT: Alignment, Classification and Tree Service (www.arb-silva.de/aligner) online server (Pruesse, Peplies & Glöckner, 2012). For this analysis, the small subunit (SSU) category was selected, and a minimum similarity identity of 0.95 was set, with ten neighbors per query sequence. Sequences below 70% identity were rejected and discarded. The least common ancestor (LCA) method was used for assignment with all five taxonomic databases ticked for classification, i.e. SILVA, Ribosomal Database Project (RDP), GreenGenes, LTP and EMBL. The SILVA database (Pruesse et al., 2007) showed more ASVs with the higher taxonomic level assignment and therefore used for subsequent analysis.

### Supplementary Figures

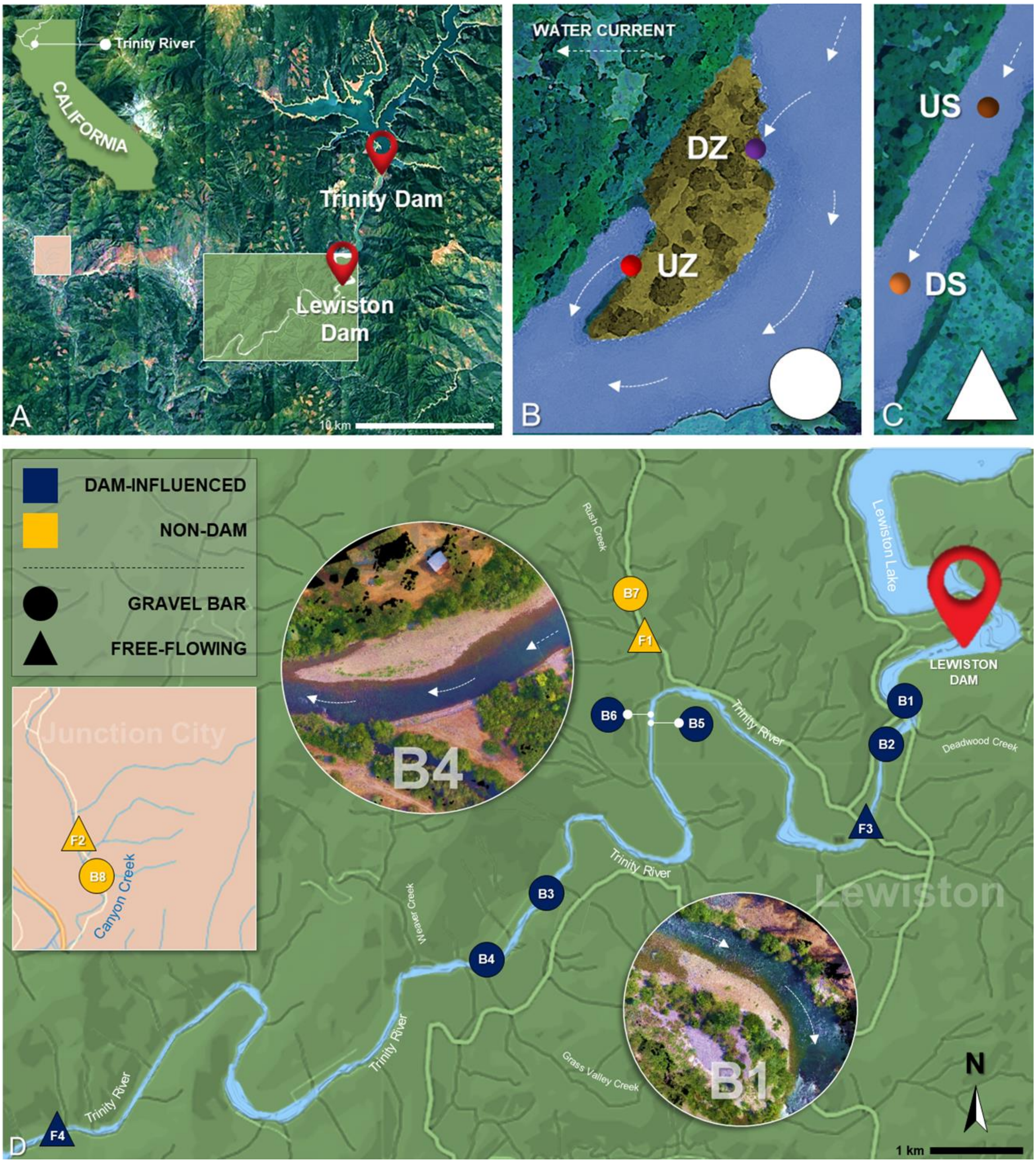


**Fig. S1** Sampling location. **a** The Trinity River and its tributaries impounded by the Trinity Dam and the Lewiston Dam in California, USA. **b** Gravel bars were sampled at the head (down-welling zone; DZ) and tail (up-welling zone; UZ) points. **c** Free-flowing sites were assessed, and samples were collected on up- (US) and downstream (DS) points. **d** In total, eight gravel bars and four free-flowing sites were assessed in the study; gravel bars B1 and B4 drone photos shown as examples. Parts of the map was generated from Google Earth Pro (version 7.3.2.5776; https://www.google.com/earth/).

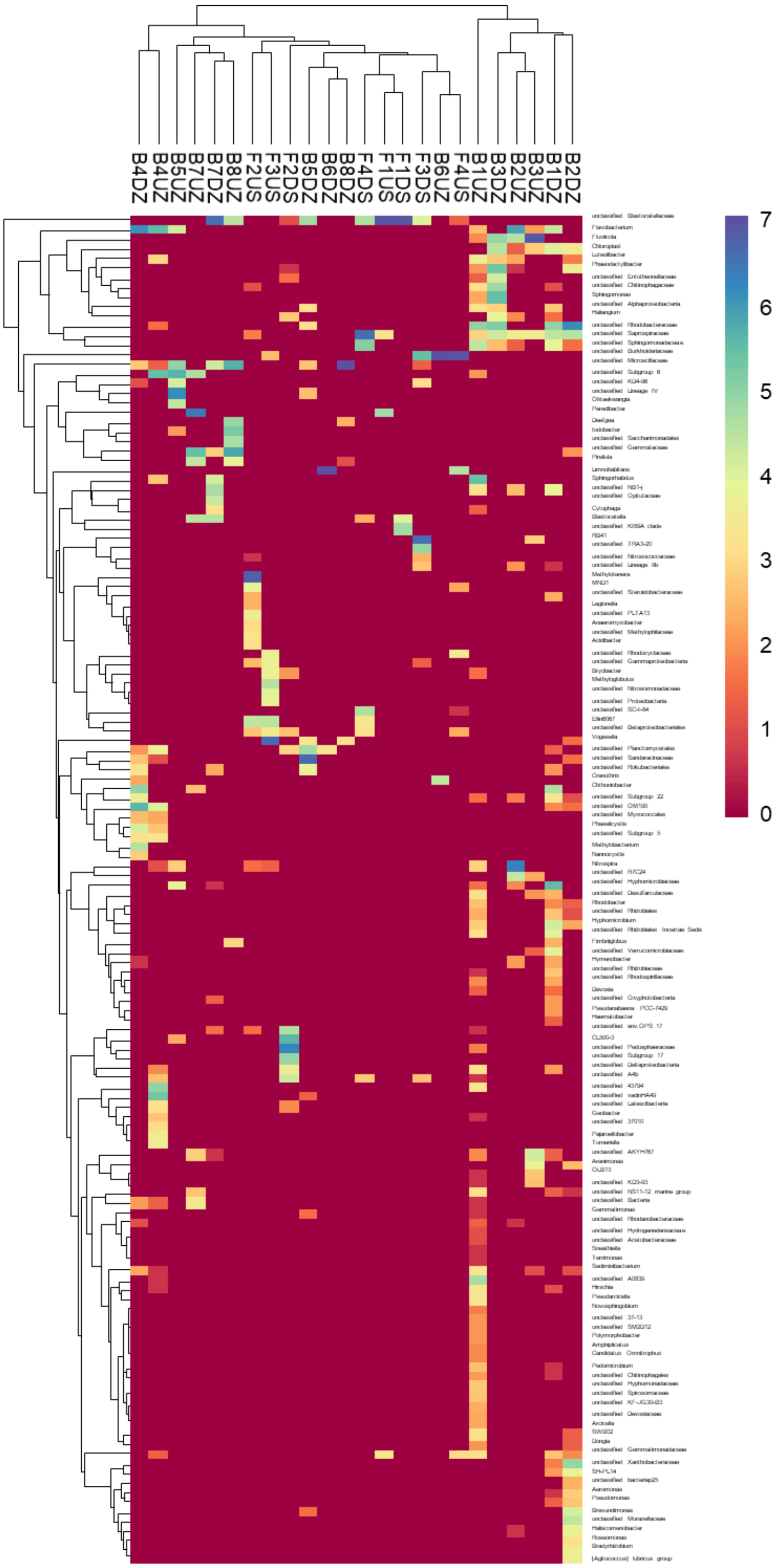


**Fig. S2** The relative abundance of the microbial community composition of the 24 sampling points at the genus-level (showing taxa with >1% read abundance). Double hierarchical cluster-based on Euclidean distance performed on rarefied and log-transformed data.

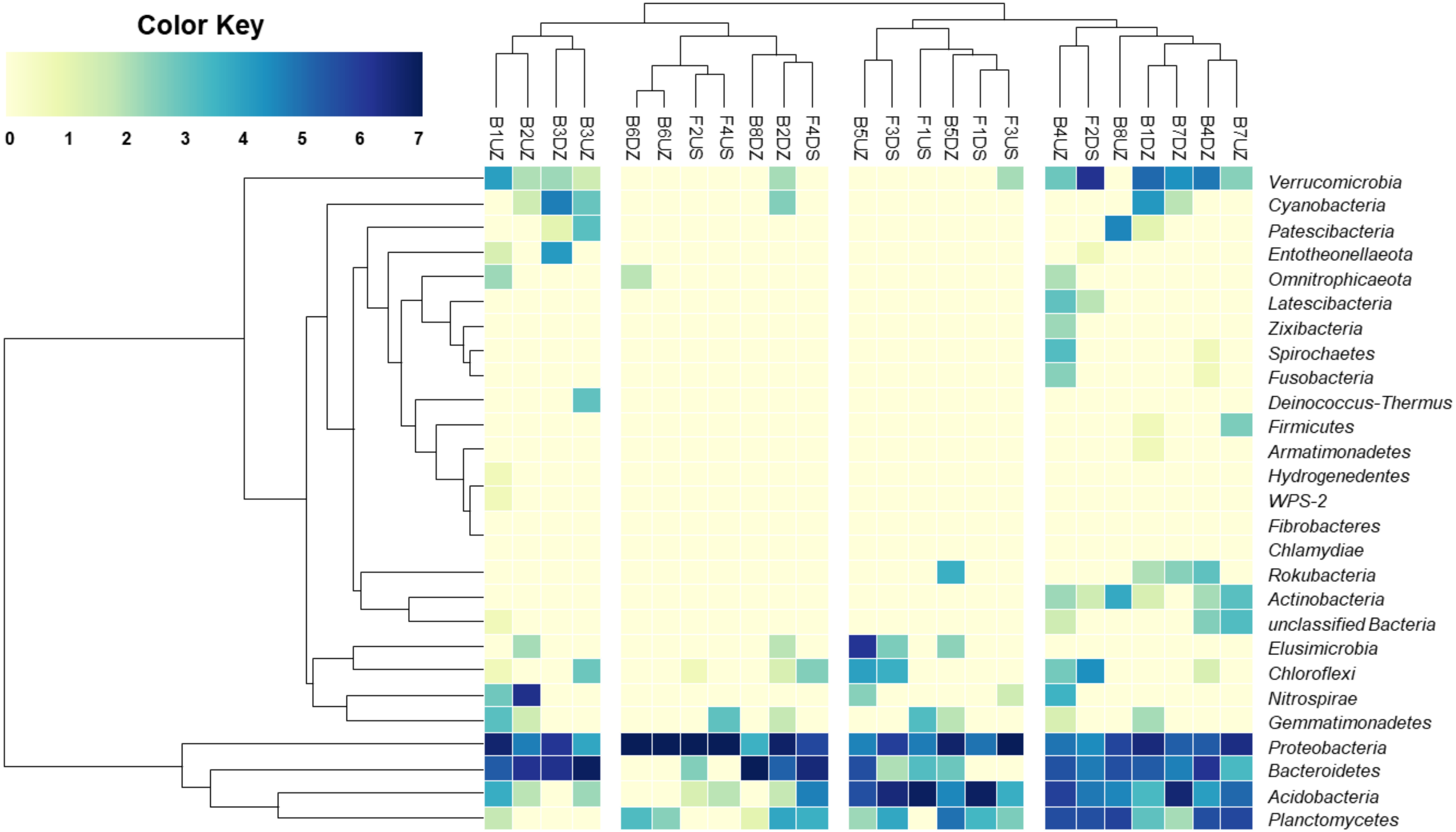


**Fig. S3** The relative abundance of the microbial community composition of the 24 sampling points at the phylum-level. Double hierarchical cluster-based on Euclidean distance performed on rarefied and log-transformed data. Columns grouped in 4 clusters.

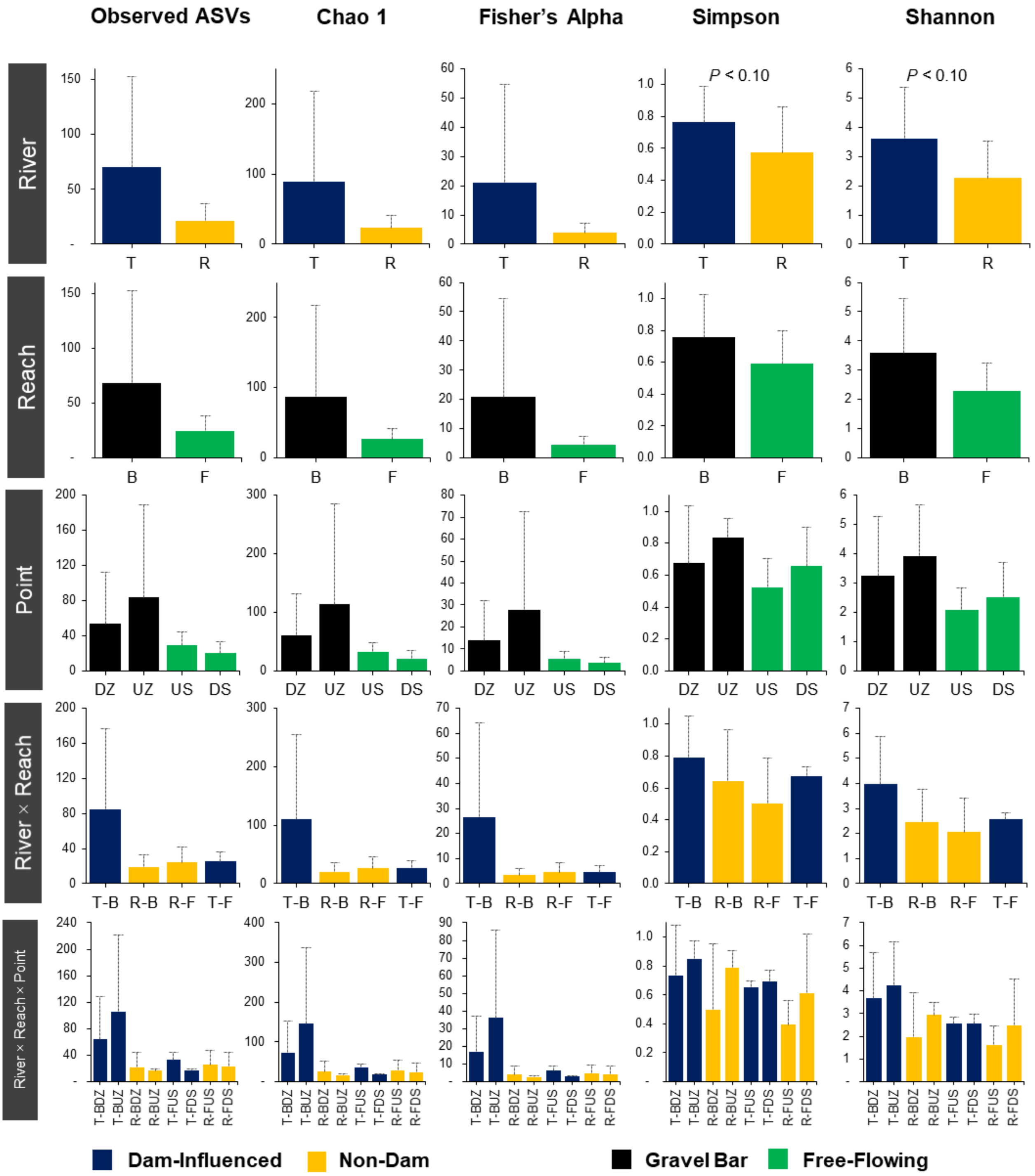


**Fig. S4** Alpha diversity estimates (mean value) for each categorical factors. Location: "T" stands for Trinity River (dam-influenced); "R" for reference creek (no dam-influence). Site: "B" for gravel bar; "F" for the free-flowing segment of the river. Point: "UZ" for up- and "DZ" for down-welling zones of the gravel bar sites; "US" for up- and "DS" for down-stream sampling points of the free-flowing river segments.

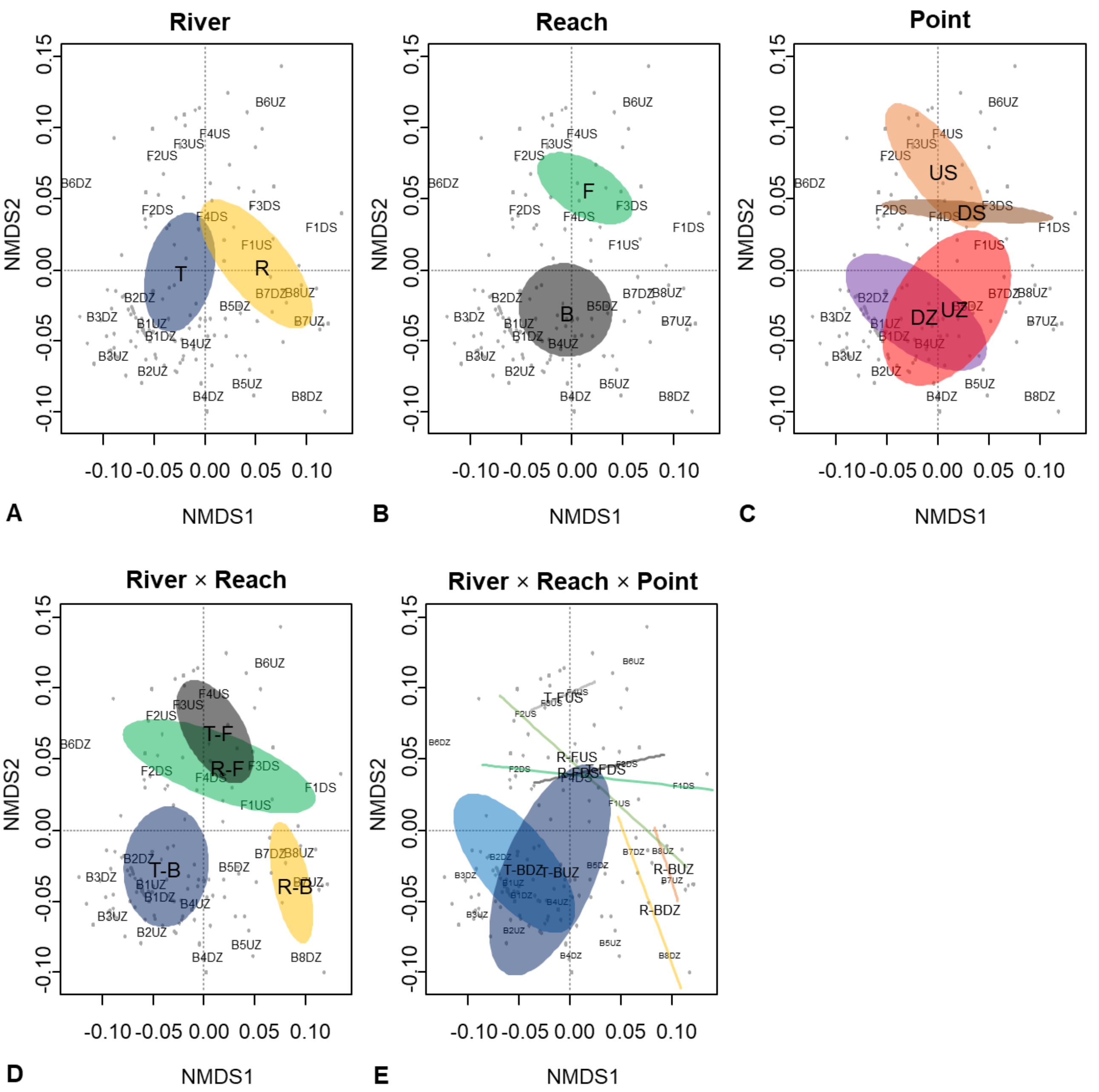


**Fig. S5** Microbial community composition at the genus-level of all 24 sampling points indicated by a non-metric multidimensional scaling plot (NMDS). Groupings based on the categorical factors are presented with ellipses (95% confidence) representing the standard error around the centroid. **a** River: “T” stands for Trinity River (dam-influenced); “R” for reference creek (no dam-influence). **b** Reach: “B” for gravel bar; “F” for the free-flowing segment of the river. **c** Point: “UZ” for up- and “DZ” for down-welling zones of the gravel bar sites; “US” for up- and “DS” for down-stream sampling points of the free-flowing river segments, and their combinations: **d** “River × Reach”, and **e** “River × Reach × Point”.

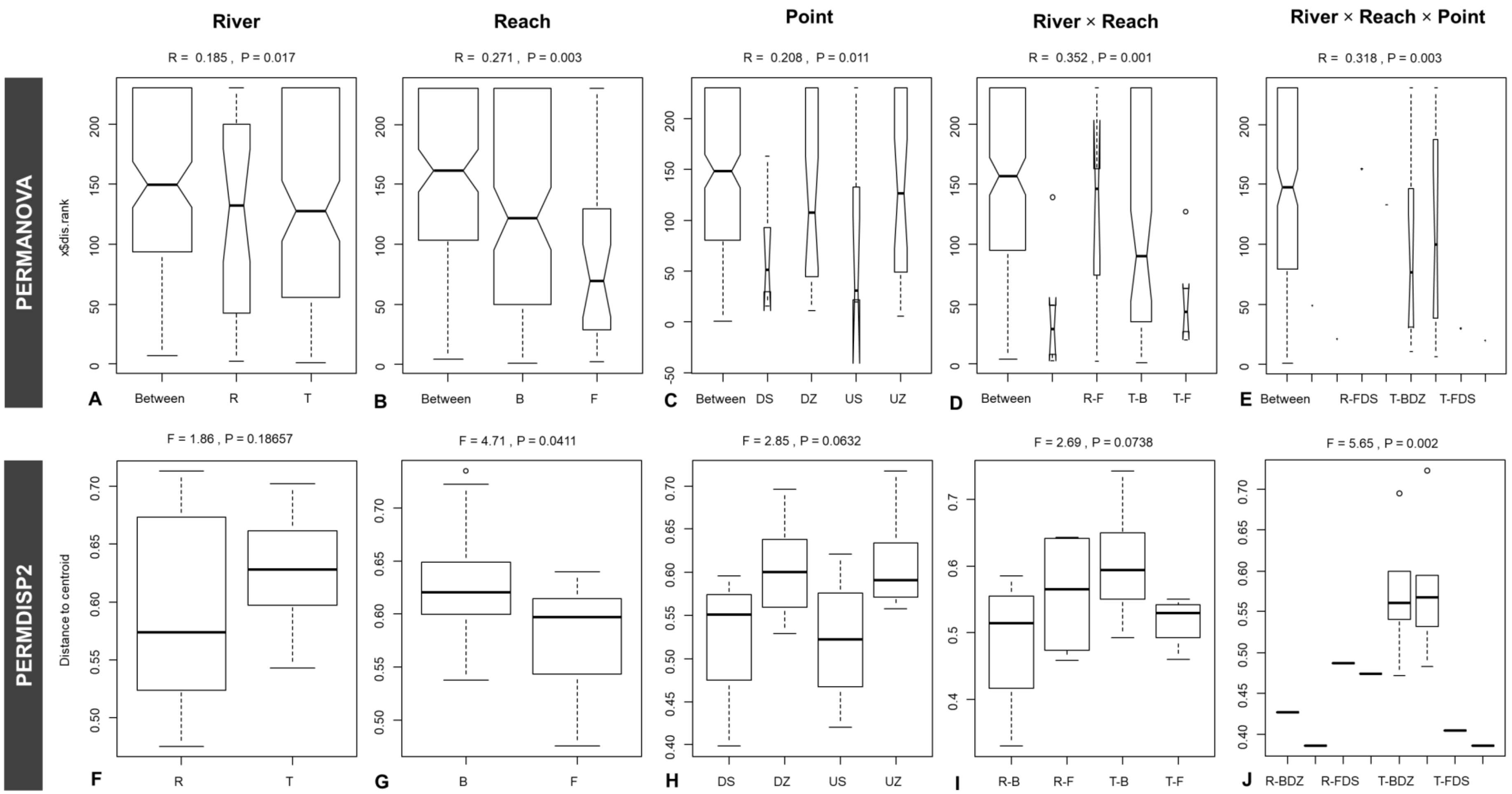


**Fig. S6** Box plots of distances to group centroids from PERMANOVA (A-E) and PERMDISP2 (F-G). Groupings based on the categorical factors. River: “T” stands for Trinity River (dam-influenced); “R” for reference creek (no dam-influence). Reach: “B” for gravel bar; “F” for the free-flowing segment of the river. Point: “UZ” for up- and “DZ” for down-welling zones of the gravel bar sites; “US” for up- and “DS” for down-stream sampling points of the free-flowing river segments.

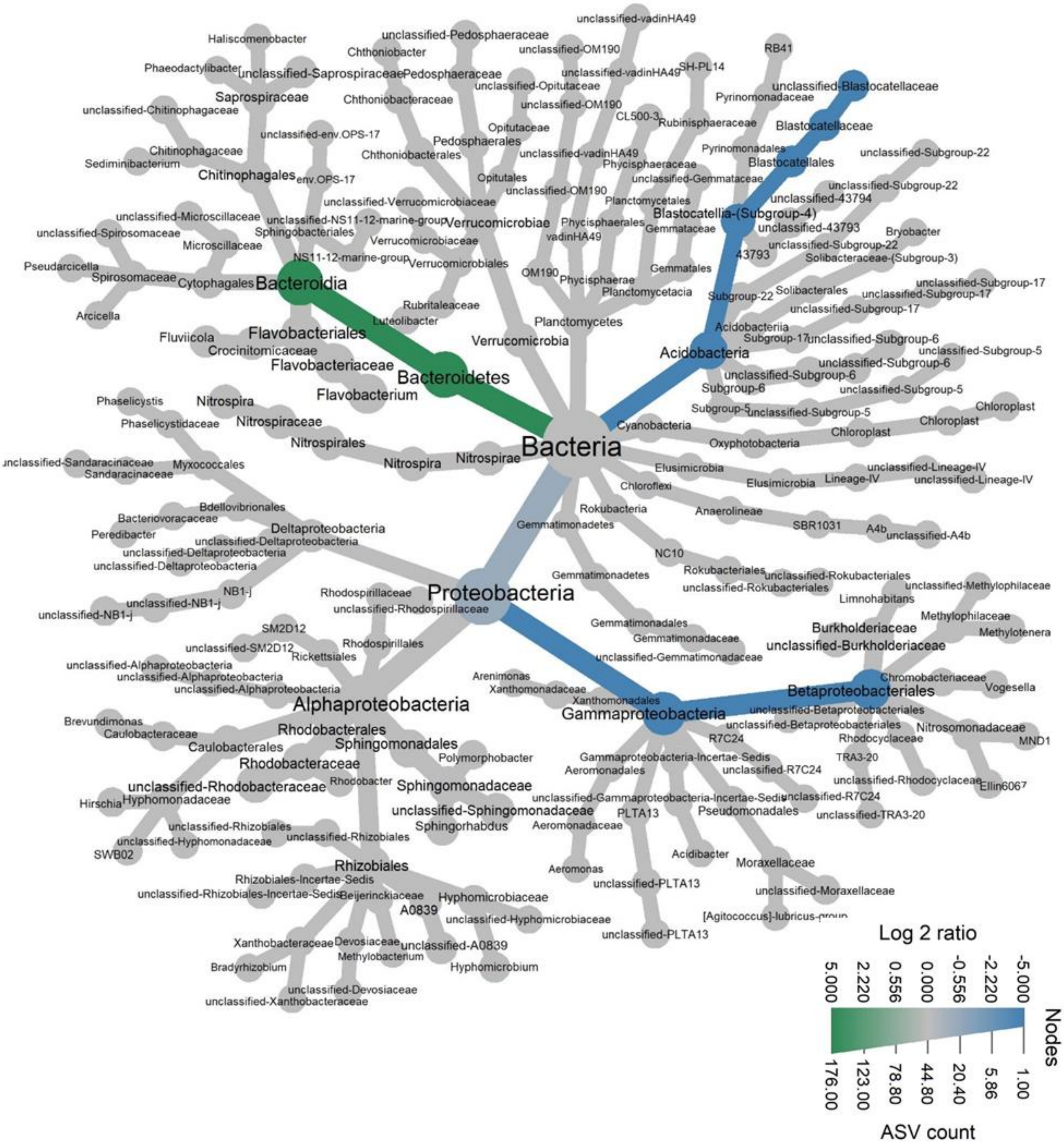


**Fig. S7** Pairwise taxonomic comparisons (ASVs with relative read abundance >0.10) between the "Reach" categories. Taxon colored in blue are relatively more abundant (in numbers of ASVs) in the gravel bars and the taxon colored green is more abundant in the free-flowing reaches. Note that this is a taxonomic tree, not a phylogenetic tree. Legend: width indicates number of ASVs at a given taxonomic rank, and color indicates relative differences in log2 (number of ASVs).

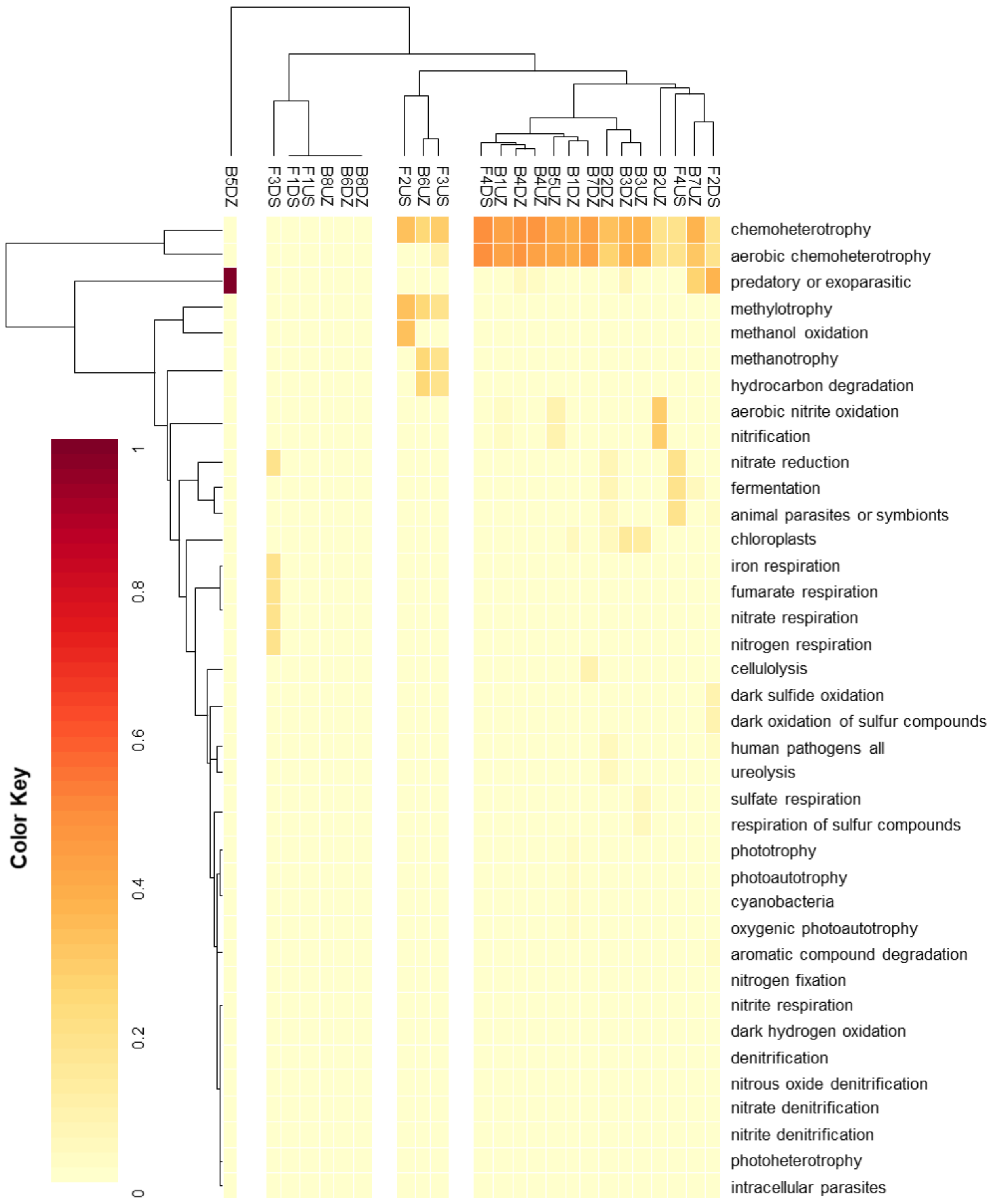


**Fig. S8** The relative abundance of the predicted functional groups of the 24 sampling points. Double hierarchical cluster-based on Euclidean distance. Columns grouped in 4 clusters.

### Supplementary Tables

**Table S1** Physico-chemical properties. “DZ” represents the down-welling and “UZ” up-welling zones of a gravel bar (B); “US” represents the up-stream and “DS” down-stream points of collection for the free-flowing segments of the rivers (F) (approx. 20-m length). “EC” stands for electric conductivity; “DO” for dissolved oxygen; “$NH_4$” for ammonium nitrogen; “AFDM” for ash-free dry mass; “TSS” for total suspended solids; and “$NO_3$” for nitrate nitrogen.

| **Sampling Site** | | **pH** | **EC** (µS/cm) | **DO** (mg/L) | **$NH_4$** (mg/L) | **AFDM** (mg/m3) | **TSS** (mg/L) | **$NO_3$** (mg/L) |
|---|---|---|---|---|---|---|---|---|
| Trinity River Gravel Bars | B1DZ | 6.90 | 82 | 10.30 | 0.03 | 200 | 0.90 | 0.04 |
| | B1UZ | 6.80 | 81 | 5.94 | 0.02 | 400 | 1.10 | 0.04 |
| | B2DZ | 7.40 | 80 | 5.99 | 0.03 | 400 | 1.30 | 0.04 |
| | B2UZ | 7.00 | 82 | 14.98 | 0.02 | 400 | 2.60 | 0.06 |
| | B3DZ | 7.10 | 85 | 5.55 | 0.02 | 1,500 | 19.50 | 0.06 |
| | B3UZ | 7.50 | 82 | 5.35 | 0.01 | 1,200 | 7.60 | 0.04 |
| | B4DZ | 7.50 | 86 | 4.71 | 0.02 | 200 | 0.90 | 0.06 |
| | B4UZ | 7.10 | 127 | 0.76 | 0.02 | 800 | 22.60 | 0.09 |
| | B5DZ | 7.90 | 85 | 6.12 | 0.01 | 400 | 0.80 | 0.06 |
| | B5UZ | 7.50 | 85 | 5.14 | 0.01 | 1,800 | 10.40 | 0.05 |
| | B6DZ | 7.10 | 85 | 7.15 | 0.01 | 200 | 1.10 | 0.06 |
| | B6UZ | 7.40 | 81 | 11.65 | 0.01 | 400 | 4.30 | 0.06 |
| Tributary Gravel Bars | B7DZ | 7.80 | 119 | 6.05 | 0.01 | 52,400 | 1.80 | 0.00 |
| | B7UZ | 7.60 | 118 | 4.74 | 0.01 | 82,100 | 2.40 | 0.00 |
| | B8DZ | 7.40 | 67 | 4.71 | 0.01 | 12,800 | 1.00 | 0.05 |
| | B8UZ | 7.80 | 65 | 4.25 | 0.01 | 25,200 | 11.40 | 0.05 |
| Tributary River Segment | F1US | 7.70 | 117 | 5.86 | 0.02 | 12,400 | 1.10 | 0.05 |
| | F1DS | 7.50 | 117 | 6.69 | 0.06 | 9,890 | 0.90 | 0.00 |
| | F2US | 7.80 | 65 | 9.47 | 0.01 | 13,100 | 1.00 | 0.00 |
| | F2DS | 7.80 | 64 | 5.34 | 0.02 | 9,290 | 0.90 | 0.00 |
| Trinity River Segment | F3US | 7.40 | 94 | 13.16 | 0.04 | 615 | 0.80 | 0.00 |
| | F3DS | 7.60 | 92 | 12.17 | 0.01 | 575 | 0.90 | 0.00 |
| | F4US | 7.50 | 92 | 14.82 | 0.03 | 33 | 0.50 | 0.00 |
| | F4DS | 7.70 | 96 | 13.01 | 0.01 | 968 | 0.70 | 0.00 |
| Variance Inflation Factor, VIF | | 1.93 | 1.63 | 2.50 | 2.05 | 2.89 | 1.74 | 3.02 |

**Table S2** PERMDISP2 analysis to test the effect of the categorical factors on the physicochemical parameters.

| **Categorical Factor** | **pH** | **EC** (µS/cm) | **DO** (mg/L) | **$NH_4$-N** (mg/L) | **AFDM** (mg/m3) | **TSS** (mg/L) | **$NO_3$-N** (mg/L) |
|---|---|---|---|---|---|---|---|
| River | 2.29 | 46.14*** | 2.08 | 0.02 | 1.4 | 1.19 | 1.28 |
| Reach | 3.86* | 0.41 | 0.11 | 2.11 | 0.18 | 14.05*** | 0.36 |
| Point | 1.08 | 0.28 | 0.89 | 0.53 | 0.41 | 1.1 | 0.37 |
| River × Reach | 2.57* | 14.75*** | 0.83 | 1.02 | 0.1 | 3.98* | 1.36 |
| River × Reach × Point | 1.36 | 5.66*** | 0.79 | 5.18** | 2.14' | 0.54 | 11.33*** |

“River” (2 groups: Trinity River – dam-influenced, Reference River – non-dam-influenced); "Reach" (2 groups: gravel bars, free-flowing segment); “Point” (4 groups: up- and down-welling, and up- and down-stream). Significance code: ‘***’ associated with a variable at $p < 0.001$, ‘**’ at $p < 0.01$, ‘*’ at $p < 0.05$ and ‘'’ at $p < 0.1$.

**Table S3** Read processing and amplicon sequence variants (ASVs) generation.

| Sampling Site | Read Abundance | | | | | Amplicon Sequence Variants | |
|---|---|---|---|---|---|---|---|
| | input | filtered | denoisedF | nonchim | reads w/ Tax. ID | ASV | ASV w/ Tax ID |
| B1DZ | 517,451 | 145,255 | 103,862 | 99,162 | 93,742 | 242 | 181 |
| B1UZ | 507,243 | 333,151 | 310,149 | 288,826 | 265,421 | 1,127 | 797 |
| B2DZ | 662,273 | 75,484 | 63,116 | 60,573 | 52,086 | 292 | 191 |
| B2UZ | 419,718 | 102,910 | 81,983 | 75,218 | 70,113 | 145 | 95 |
| B3DZ | 400,620 | 30,118 | 12,929 | 12,676 | 4,900 | 46 | 26 |
| B3UZ | 258,267 | 45,165 | 36,281 | 34,659 | 32,129 | 128 | 71 |
| B4DZ | 268,055 | 69,813 | 41,532 | 41,397 | 35,779 | 57 | 46 |
| B4UZ | 212,876 | 66,974 | 54,937 | 51,462 | 45,474 | 359 | 237 |
| B5DZ | 239,884 | 39,249 | 4,427 | 4,427 | 3,697 | 48 | 24 |
| B5UZ | 206,662 | 17,234 | 7,580 | 7,580 | 7,438 | 18 | 14 |
| B6DZ | 331,289 | 14,947 | 2,939 | 2,939 | 1,257 | 5 | 3 |
| B6UZ | 275,384 | 12,783 | 4,927 | 4,927 | 3,731 | 18 | 8 |
| B7DZ | 115,997 | 28,063 | 20,999 | 18,259 | 17,993 | 45 | 41 |
| B7UZ | 43,384 | 8,880 | 4,765 | 4,765 | 4,237 | 36 | 19 |
| B8DZ | 107,294 | 18,865 | 4,636 | 4,636 | 4,636 | 6 | 6 |
| B8UZ | 126,781 | 11,300 | 5,793 | 5,615 | 5,519 | 18 | 15 |
| F1US | 235,521 | 10,567 | 4,957 | 4,957 | 4,929 | 11 | 10 |
| F1DS | 43,038 | 16,200 | 5,444 | 5,390 | 5,238 | 13 | 9 |
| F2US | 279,684 | 92,581 | 79,151 | 78,276 | 77,656 | 80 | 73 |
| F2DS | 114,071 | 25,857 | 14,572 | 14,496 | 14,104 | 53 | 42 |
| F3US | 408,804 | 33,061 | 25,618 | 23,414 | 22,555 | 53 | 45 |
| F3DS | 138,232 | 21,716 | 16,300 | 15,502 | 14,902 | 26 | 20 |
| F4US | 338,380 | 125,162 | 113,533 | 112,457 | 112,197 | 52 | 50 |
| F4DS | 188,203 | 10,511 | 5,907 | 5,850 | 5,629 | 23 | 17 |
| **Total** | **6,439,111** | **1,355,846** | **1,026,337** | **977,463** | **905,362** | **2,855** | **1,998** |

**Table S4** Richness and diversity estimates of the 24 sampling points.

| Site | Observed ASVs | Chao1 | Fisher's alpha | Simpson | Shannon |
|---|---|---|---|---|---|
| B1DZ | 149 | 179.8750 | 43.9939 | 0.9498 | 5.7676 |
| B1UZ | 307 | 508.5000 | 129.4823 | 0.9694 | 6.8123 |
| B2DZ | 143 | 166.3333 | 41.5421 | 0.9006 | 5.2475 |
| B2UZ | 73 | 77.3333 | 16.8848 | 0.9113 | 4.4240 |
| B3DZ | 26 | 26.0000 | 4.6380 | 0.9143 | 3.8848 |
| B3UZ | 59 | 66.0909 | 12.8433 | 0.7838 | 3.2798 |
| B4DZ | 40 | 43.3333 | 7.8756 | 0.9159 | 4.2016 |
| B4UZ | 174 | 203.2162 | 54.7930 | 0.9671 | 6.1077 |
| B5DZ | 24 | 24.0000 | 4.2085 | 0.6725 | 2.7176 |
| B5UZ | 14 | 14.0000 | 2.2057 | 0.8237 | 3.0759 |
| B6DZ | 3 | 3.0000 | 0.3688 | 0.0528 | 0.1976 |
| B6UZ | 8 | 8.0000 | 1.1421 | 0.6488 | 1.8535 |
| B7DZ | 38 | 43.1429 | 7.3898 | 0.8194 | 3.3623 |
| B7UZ | 18 | 18.0000 | 2.9761 | 0.7034 | 2.5688 |
| B8DZ | 6 | 6.0000 | 0.8176 | 0.1782 | 0.5817 |
| B8UZ | 15 | 15.0000 | 2.3941 | 0.8743 | 3.3159 |
| F1US | 10 | 10.0000 | 1.4829 | 0.2792 | 1.0338 |
| F1DS | 8 | 8.0000 | 1.1421 | 0.3263 | 1.0462 |
| F2US | 41 | 46.6250 | 8.1213 | 0.5123 | 2.2159 |
| F2DS | 38 | 40.0000 | 7.3898 | 0.9017 | 3.9183 |
| F3US | 41 | 41.5000 | 8.1213 | 0.6234 | 2.7603 |
| F3DS | 19 | 19.0000 | 3.1754 | 0.7506 | 2.8385 |
| F4US | 25 | 32.0000 | 4.4221 | 0.6864 | 2.3474 |
| F4DS | 16 | 16.0000 | 2.5854 | 0.6411 | 2.2697 |

**Table S5** Multivariate pattern analysis for indicator taxa. Association function: IndVal.g. Significance code: ‘***’ associated with a variable at $p < 0.001$, ‘**’ at $p < 0.01$, ‘*’ at $p < 0.05$ and ‘’’ at $p < 0.1$.

Total number of species: 325
Selected number of species: 16
Number of species associated to 1 group: 14
Number of species associated to 2 groups: 1
Number of species associated to 3 groups: 1

List of species associated to each combination:

| **Trinity Gravel Bars (T-B)** | *stat* | *p.value* | |
|---|---|---|---|
| *Flavobacterium* | 0.764 | 0.0169 | * |
| unclassified Rhodobacteraceae | 0.707 | 0.0655 | ' |
| *Sediminibacterium* | 0.707 | 0.0681 | ' |

| **Reference Gravel Bars (R-B)** | *stat* | *p.value* | |
|---|---|---|---|
| *Pirellula* | 0.866 | 0.00603 | ** |
| unclassified Gemmataceae | 0.842 | 0.00328 | ** |
| unclassified Microscillaceae | 0.753 | 0.02377 | * |
| *Deefgea* | 0.707 | 0.06458 | ' |
| unclassified Sporichthyaceae | 0.66 | 0.07012 | ' |

| **Trinity Free-flowing Site (T-F)** | *stat* | *p.value* | |
|---|---|---|---|
| unclassified Burkholderiaceae | 0.806 | 0.0131 | * |
| unclassified Gammaproteobacteria | 0.71 | 0.0212 | * |
| SM1A02 | 0.707 | 0.0667 | ' |
| *Haemophilus* | 0.707 | 0.0666 | ' |
| unclassified Rhodocyclaceae | 0.672 | 0.0666 | ' |

| **Reference Free-flowing Site (R-F)** | *stat* | *p.value* | |
|---|---|---|---|
| *Acinetobacter* | 0.707 | 0.064 | ' |

| **T-F and R-F** (Free-flowing Reach) | *stat* | *p.value* | |
|---|---|---|---|
| unclassified Betaproteobacteriales | 0.791 | 0.0066 | ** |

| **R-B, T-F and R-F** | *stat* | *p.value* | |
|---|---|---|---|
| unclassified Blastocatellaceae | 0.799 | 0.0244 | * |

### References

Gaeuman, D. (2014). High-flow gravel injection for constructing designed in-channel features. *River Research and Applications*, *30*(6), 685-706.

Pruesse, E., Quast, C., Knittel, K., Fuchs, B. M., Ludwig, W., Peplies, J., & Glöckner, F. O. (2007). SILVA: a comprehensive online resource for quality checked and aligned ribosomal RNA sequence data compatible with ARB. Nucleic acids research, 35(21), 7188-7196.Sackett, J. D., Shope, C. L., Bruckner, J. C., Wallace, J., Cooper, C. A., & Moser, D. P. (2019). Microbial Community Structure and Metabolic Potential of the Hyporheic Zone of a Large Mid-Stream Channel Bar. *Geomicrobiology Journal*, 1-12.

Pruesse, E., Peplies, J., & Glöckner, F. O. (2012). SINA: accurate high-throughput multiple sequence alignment of ribosomal RNA genes. *Bioinformatics*, 28(14), 1823-1829.

Ock, G., Gaeuman, D., McSloy, J., & Kondolf, G. M. (2015). Ecological functions of restored gravel bars, the Trinity River, California. *Ecological engineering*, *83*, 49-60.
